# SCLC_CellMiner: Integrated Genomics and Therapeutics Predictors of Small Cell Lung Cancer Cell Lines based on their genomic signatures

**DOI:** 10.1101/2020.03.09.980623

**Authors:** Camille Tlemsani, Lorinc Pongor, Luc Girard, Nitin Roper, Fathi Elloumi, Sudhir Varma, Augustin Luna, Vinodh N. Rajapakse, Robin Sebastian, Kurt W. Kohn, Julia Krushkal, Mirit Aladjem, Beverly A. Teicher, Paul S. Meltzer, William C. Reinhold, John D. Minna, Anish Thomas, Yves Pommier

## Abstract

Model systems are necessary to understand the biology of SCLC and develop new therapies against this recalcitrant disease. Here we provide the first online resource, CellMiner-SCLC (https://discover.nci.nih.gov/SclcCellMinerCDB) incorporating 118 individual SCLC cell lines and extensive omics and drug sensitivity datasets, including high resolution methylome performed for the purpose of the current study. We demonstrate the reproducibility of the cell lines and genomic data across the CCLE, GDSC, CTRP, NCI and UTSW datasets. We validate the SCLC classification based on four master transcription factors: NEUROD1, ASCL1, POU2F3 and YAP1 (NAPY classification) and show transcription networks connecting each them with their downstream and upstream regulators as well as with the NOTCH and HIPPO pathways and the MYC genes (MYC, MYCL1 and MYCN). We find that each of the 4 subsets express specific surface markers for antibody-targeted therapies. The SCLC-Y cell lines differ from the other subsets by expressing the NOTCH pathway and the antigen-presenting machinery (APM), and responding to mTOR and AKT inhibitors. Our analyses suggest the potential value of NOTCH activators, YAP1 inhibitors and immune checkpoint inhibitors in SCLC-Y tumors that can now be independently validated.

**Graphical Abstract:** 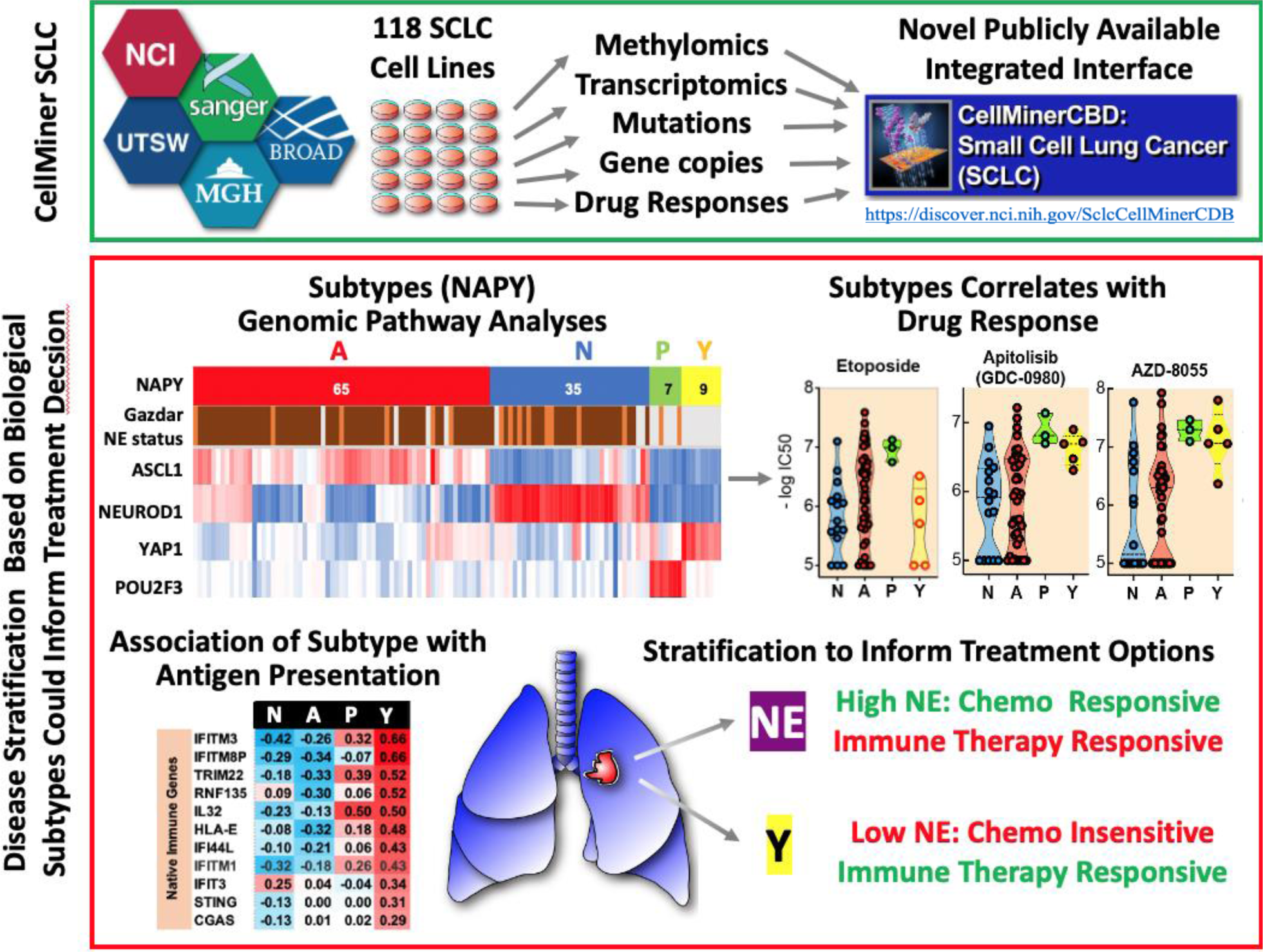

**Highlights:**

- SCLC-CellMiner provides the most extensive SCLC resource in terms of number of cell lines (118 cell lines), extensive omics data (exome, microarray, RNA-seq, copy number, methylomes and microRNA) and drug sensitivity testing.
- We find evidence of distinct epigenetic profile of SCLC cell lines (global hypomethylation and histone gene methylation), which is consistent with their plasticity.
- Transcriptome analyses demonstrate the coherent transcriptional networks associated with the 4 main genomic subgroups (NEUROD1, ASCL1, POU2F3 & YAP1 = NAPY classification) and their connection with the NOTCH and HIPPO signaling pathways.
- SCLC-CellMiner provides a conceptual framework for the selection of therapies for SCLC in a personalized fashion allowing putative biomarkers according molecular classifications and molecular characteristics.
- SCLC-Y cell lines differ from the other cancer cell lines; their transcriptome resemble NSCLC cell lines. YAP1 cell lines while being the most resistant to standard of care treatments (etoposide, cisplatin and topotecan) respond to mTOR and AKT inhibitors and present native immune predisposition suggesting sensitivity to immune checkpoint inhibitors.

## Introduction

Lung cancer is the leading cause of cancer death worldwide. Although small cell lung cancer (SCLC) represents only 15% of all lung cancers, it accounts for more than 30,000 cases in the US alone and has the most aggressive clinical course with most patients presenting with widely metastatic disease and a median survival of 10-12 months (Wang et al., 2017). The diagnosis of SCLC is based on histological features including dense sheets of small cells with scant cytoplasm, ill-defined borders and nuclei with finely granular chromatin lacking prominent nucleoli (Gazdar et al., 2017; Hann et al., 2019; Rudin et al., 2019). Immunohistochemistry shows high Ki-67, consistent with rapid cellular proliferation generally driven by high MYC oncogenic expression together with tumor suppressor RB1 and TP53 inactivation (Gazdar et al., 2017). Unlike the increasingly personalized treatment approaches for non-small cell lung cancer (NSCLC), SCLC is currently treated as a homogeneous disease (Hann et al., 2019; Rudin et al., 2019; Thomas and Pommier, 2016). The typical low life expectancy for a patient diagnosed with SCLC and the options for therapy (platinum-etoposide combination as first line therapy and topotecan at relapse) remain limited, causing the National Cancer Institute (NCI) to categorize SCLC as a “recalcitrant” cancer.

Most SCLC tumors are characterized by their neuroendocrine differentiation, which can be histologically visualized using a panel of markers including synaptophysin (SYP), chromogranin A (CHGA), NCAm1 and insulinoma-associated protein 1 (INSM1) (Gazdar et al., 2017; Hann et al., 2019; McColl et al., 2017). Yet, a smaller subset of SCLC is negative for the standard neuroendocrine markers (Gazdar et al., 2017; Guinee et al., 1994; Hann et al., 2019; McColl et al., 2017). Hence, SCLCs have been historically defined as “classic” (neuroendocrine: NE) or “variant” (non-neuroendocrine: non-NE) (Gazdar et al., 2017; Gazdar et al., 1985; Rudin et al., 2019). Ongoing efforts are designed to categorize the molecular subtypes of SCLCs (Gazdar et al., 2017; George et al., 2015; McColl et al., 2017; Rudin et al., 2019) and to rationalize novel therapeutic approaches based on molecular genomic characteristics of the disease (Gardner et al., 2017; McColl et al., 2017; Thomas and Pommier, 2016).

To discriminate NE and non-NE SCLC, Gazdar *et al*, proposed a classification based on the expression of 50 genes including *ASCL1* (achaete-scute homolog 1) and *NEUROD1* (neurogenic differentiation factor 1), which are key transcription factors binding to E-box-containing promoter consensus core sequences 5’-CANNTG. ASCL1 and NEUROD1 drive the maturation of neuroendocrine cells of the lung (Borges et al., 1997; Ito et al., 2000; Neptune et al., 2008) and are highly expressed in NE SCLCs (Zhang et al., 2018). A consensus nomenclature for four molecular subtypes has been recently proposed based on differential expression of two additional transcription factors, YAP1 (Yes-Associated Protein 1) and POU2F3 (POU class 2 homeodomain box 3) for the non-NE SCLC subtype (Rudin et al., 2019). *POU2F3* encodes a member of the POU domain family of transcription factors normally expressed in rare chemosensory cells of the normal lung epithelium (tuft cells) and of the gastrointestinal track (Huang et al., 2018). Selective expression of POU2F3 was identified recently by CRISPR screening in a subset of SCLC cells that lack NE features (Huang et al., 2018). YAP1, a key mediator of the Hippo signaling pathway, was discovered as being reciprocally expressed relative to the neuroendocrine transcription factor INSM1 (McColl et al., 2017). Hence, it has now been proposed to classify SCLCs into 4 groups based on the expression of *NEUROD1, ASCL1, POU2F3* and *YAP1* (Rudin et al., 2019). For short, we will refer to this classification as “NAPY” ((N=*NEUROD1*, A=*ASCL1*, P=*POU2F3* and Y= *YAP1*) in the present study.

Genomic initiatives have accelerated the pace of discovery for many cancers (Cancer Genome Atlas Research, 2012, 2014). Unfortunately, the TCGA was not extended to SCLC because of a lack of readily accessible and adequate tumor tissue, as most patients are diagnosed with SCLC by fine-needle aspiration, while surgically resected specimens are relatively rare. Further underscoring this issue, comprehensive genomic and transcriptomic data is available only for less than 250 SCLC tumors to date. Nevertheless, SCLC research has benefited from the systematic collection of a large number of tumor cell lines; most of them developed at the US National Cancer Institute (NCI) in the NCI-VA/NCI-Navy Medical Oncology Branches (Carney et al., 1985; Gazdar et al., 1985). This collection has been distributed widely, and detailed genetic and pharmacological annotation available from several groups including the NCI, the Broad-MIT and the Sanger/MGH (Barretina et al., 2012; Garnett et al., 2012; Polley et al., 2016). Yet, in spite of large number of cell lines and drugs profiled (Figure 1), the data are accessible only from different platforms making it challenging to systematically translate and integrate genomic data into knowledge of SCLC tumor biology and therapeutic possibilities. Additionally, a number of SCLC cell lines generated by the Minna-Gazdar group at UT Southwestern Medical Center (McMillan et al., 2018) had not been integrated in the preexisting NCI, Broad Institute (CCLE/CTRP) and Sanger-Massachusetts General Hospital (GDSC) databases.

**Figure 1.**
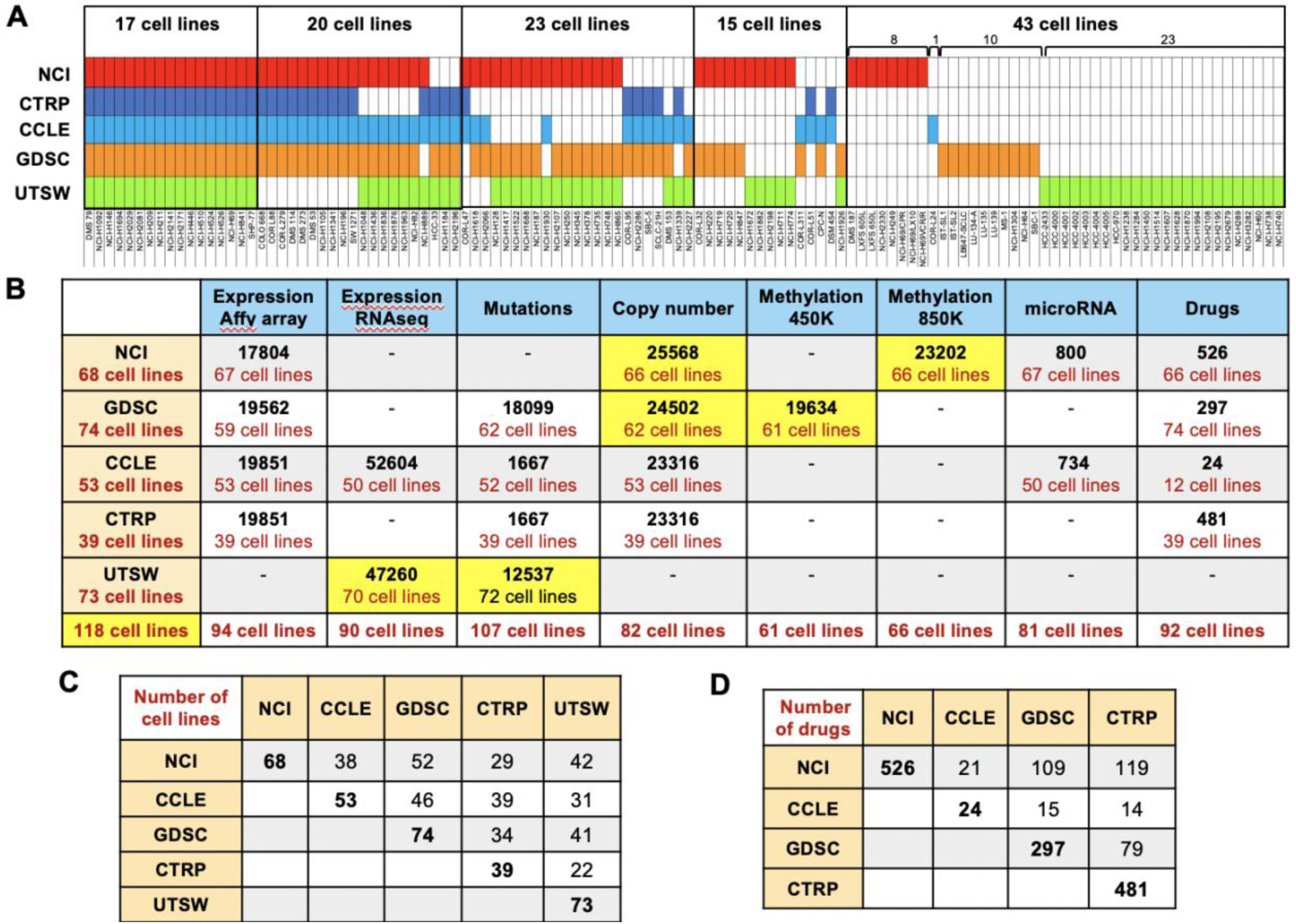
Summary of the data included in SCLC-CellMiner and resources. (**A**) Cell line overlap between the five data sources. Each colored box represents one cell line. The cell lines in red are from the NCI database (N = 68), in dark blue from CTRP (N = 39), in light blue from CCLE (N = 53), in orange from GDSC (N = 74) and in green from UTSW (N = 73). Cell line details are provided in Table S1. (**B**) Summary of the genomic and drug activities data for the five data sources in SCLC CellMinerCDB (https://discover.nci.nih.gov/SclcCellMinerCDB/). The number of SCLC cell lines for datasets and sources are indicated. For microarray, mutations, copy number and methylation data, the numbers indicate the number of genes. For RNA-seq data, the numbers indicate the number of transcripts. The bottom row show the total number of cell lines (N = 118) integrated in SCLC CellMinerCDB. New data analyses performed and made available are highlighted in yellow. (**C**) Cell line overlap between data sources. Details of the cell line overlap are provided in Table S2. (**D**) Drug overlap between data sources.

To substantially extend our understanding of the genomic features of SCLC, we performed genome-wide DNA methylation at single-base resolution by IIllumina Methylation 850k analysis on the NCI set of 68 SCLC cell lines and whole genome RNA-seq for 72 cell lines of the UTSW set. We also integrated these data in a global drug and genomic database (SCLC_Global) encompassing a total of 118 individual SCLC cell lines. This enabled us to enrich for the least represented SCLC subtypes, which are the non-NE YAP1 and POU2F3 subtypes and to further analyze the genomic and drug response characteristics of the YAP1 subgroup compared to the classical neuroendocrine NEUROD1 and ASCL1 subtypes of SCLC. The integrated data are available from the web-based tool, which we refer to as SCLC-CellMinerCDB (https://discover.nci.nih.gov/SclcCellMinerCDB/).

## Results

### SCLC-CellMinerCDB Resource

SCLC-CellMinerCDB integrates genomic and drug activity data for total of 118 molecularly characterized SCLC cell lines (Figure 1) including 68 from the NCI (Polley et al., 2016), 74 from the GDSC (Garnett et al., 2012), 53 from the CCLE, 39 from the CTRP (Barretina et al., 2012) and 73 from UT Southwestern (UTSW) (Gazdar et al., 2010). Details for each cell line (source of the cell lines with patient characteristics and main genomic features and classification) is provided in Supplemental Table S1. Among those 118 SCLC cell lines, 17 (14%) are in all five data sources, 20 (17%) are in four data sources, 23 (20%) in three data sources, 15 (13%) in two data sources while 43 (36%) are present in only one data source (Figure 1A and Supplemental Table S2).

Our integrated resource includes new data obtained by performing high resolution whole genome methylome and copy number analyses for 66 cell lines as well as whole genome-level transcriptome by RNA-seq for 72 cell lines. Data first made available here are highlighted with yellow background in Figure 1B. SCLC-CellMinerCDB also makes accessible whole exome mutation data for 12,537 genes across 72 cell lines of the UTSW SCLC database in addition to the previously released whole exome sequencing data for 52 cell lines from CCLE and 62 cell lines from GSDC.

The range of tested clinical drugs and investigational compounds in each dataset and across data sources is summarized in Figure 1D. The NCI database provides the largest number of tested compounds (N = 526), followed by the CTRP (N = 481), GDSC (N = 297) and CCLE (N = 224). The overlap between tested compound across the data sources is also shown in Figure 1D.

SCLC_CellMiner allows multiple analyses listed in Table 1. They include confirming cell line consistency and identity across datasets, drug activity reproducibility across datasets, determinants of gene expression (based on DNA copy number, promoter methylation and microRNA expression), exploration and validation of genomic networks, classification of the cell lines based on metadata such as the NAPY, epithelial mesenchymal (EMT) and antigen presenting machinery (APM) scores and the validation and discovery of drug response determinants.

**Table 1:**
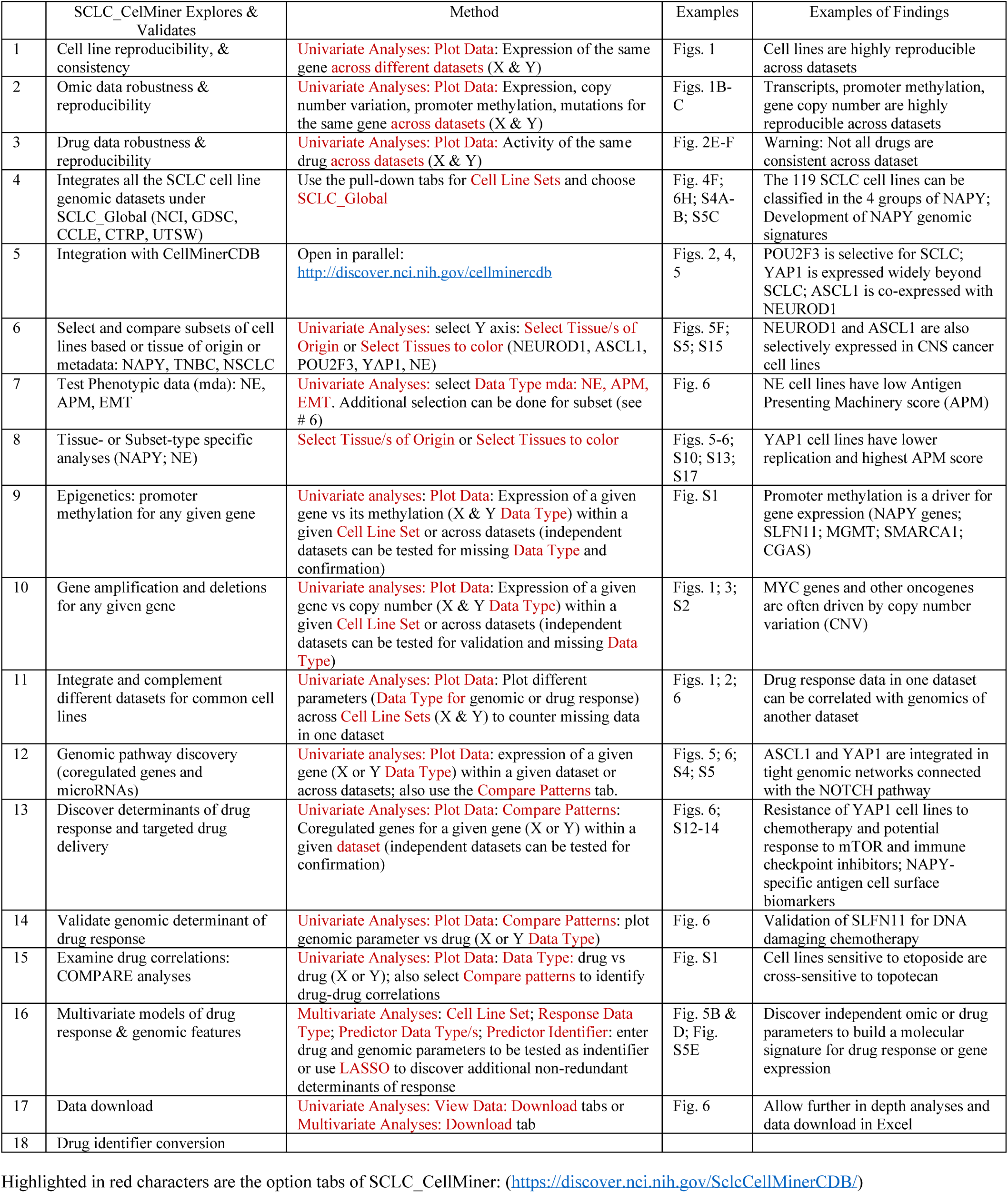
Examples of SCLC_CellMiner capabilities:

### Data Validation, Cross-Database (CDB) Analyses and CellMinerCDB Univariate Analyses

Cross comparison for matched cell lines between databases was used to validate the new NCI-SCLC methylome (850K Illumina array) by comparison with the published SCLC data of GDSC (450K array) (Rajapakse et al., 2018). The comparison yielded remarkably high overall correlation with a median of 0.92 for 7,246 common genes with with wide expression range for the 43 common cell lines (Figure 2A). Cross-correlation of the new RNA-seq data from UTSW with other gene expression data (microarray and RNA-seq) were also highly significant albeit with lower median correlations (Figure 2A). These data demonstrate the high reproducibility of the new data (NCI methylome and UTSW RNA-seq) (McMillan et al., 2018) across independent databases and the similarity of cell lines grown at different institutions and analyzed independently with different technical platforms (RNA-seq vs microarray, 850k vs 450k methylome arrays).

**Figure 2.**
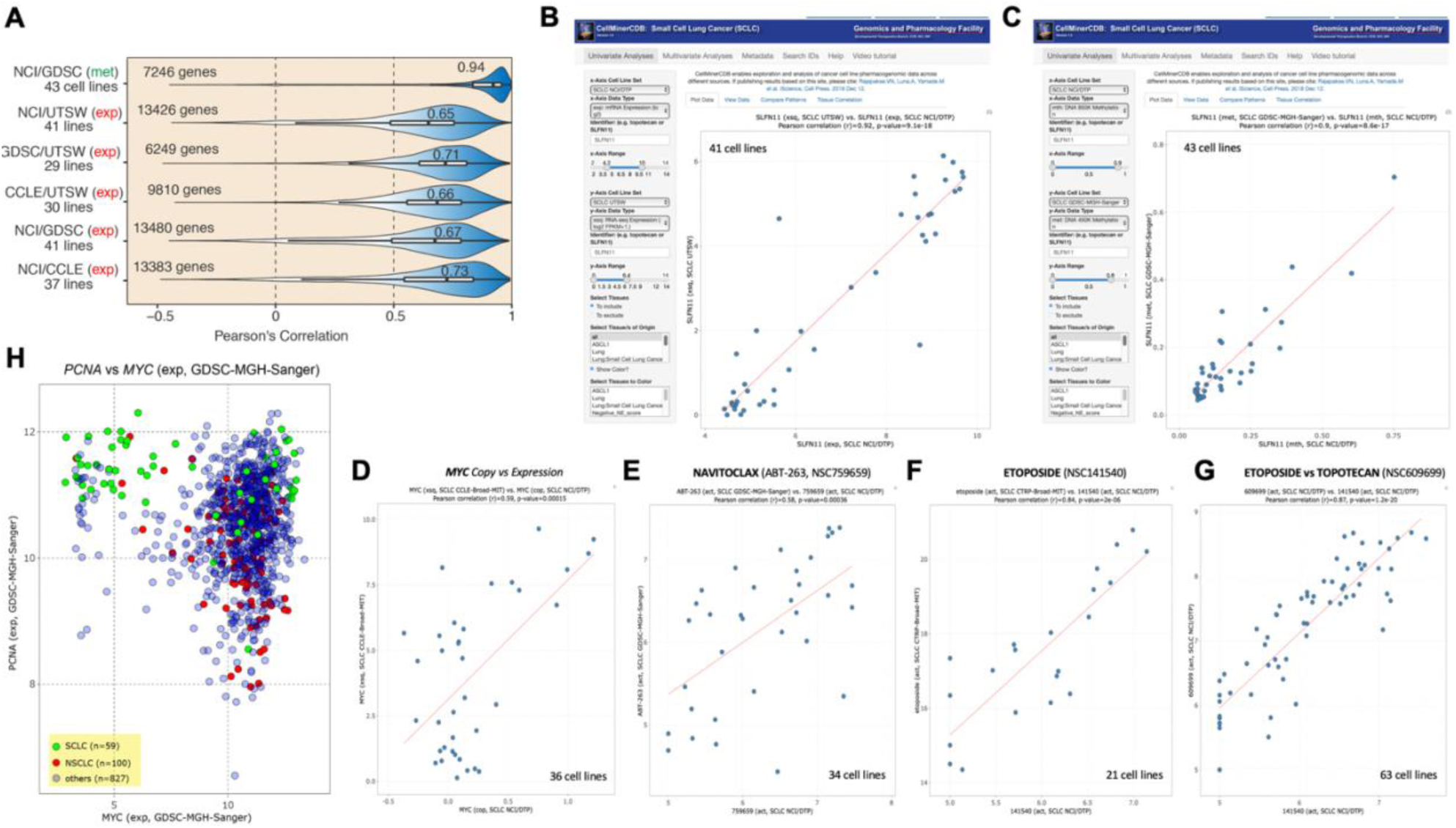
Validation and reproducibility of the SCLC-CellMiner data with snapshots illustrating representative outputs of SCLC-CellMiner (https://discover.nci.nih.gov/SclcCellMinerCDB/) (**A**) Reproducibility between data sources. The figure represents the expression and methylation Pearson correlations between the indicated data sources for matched cell lines (see Figure 1). The median of expression Pearson correlation is 0.65, 0.67, 0.73, 0.66 and 0.71 for NCI /UTSW, NCI/GDSC, NCI/CCLE, UTSW/CCLE, and UTSW/GDSC, respectively. The median of methylation Pearson correlation between NCI and GDSC data sources is 0.94. (**B**) Snapshot from SCLC-CellMiner showing the reproducibility of *SLFN11* gene expression across the 41 common cell lines independently of the methods used to measure *SLFN11* expression (AffyArray for NCI/DTP on the x-axis vs RNA-Seq for UTSW). Each dot is a different cell line, which can be identified by moving the cursor to the dot on the CellMiner website. The data can also be readily displayed in tabular form and downloaded in tab-delimited format by clicking on the “View Data” tab to the right of the default “Plot Data” tab (see upper section of Figures 2B & C). (**C**) Snapshot from SCLC-CellMiner showing the reproducibility of *SLFN11* promoter methylation across the 43 common cell lines independently of the methods used to measure *SLFN11* expression (850 k Illumina Infinium MethylationEPIC BeadChip array for NCI/DTP on the x-axis vs Illumina HumanMethylation 450K BeadChip array for GDSC). (**D**) SCLC-CellMiner demonstrates the highly significant correlation between *MYC* DNA copy number (new data derived from the 850 K AffyArray methylome of the NCI-SCLC cell lines and *MYC* expression (data from CCLE) for the 36 common cell lines. (**E**-**G**) Examples (image snapshots from SCLC-CellMiner) of drug activity correlations across databases for the indicated drugs and the common cell lines) (**H**) High proliferation signature of SCLC cell lines based on high *PCNA* and *MYC* expression. Note that SCLC (green) overexpress PCNA but fall into two groups with respect to MYC (high and low). The image was obtained through CellMinerCDB with the GDSC database (http://discover.nci.nih.gov/cellminercdb).

Measurement reproducibility across pharmacogenomic datasets can instantly be performed and displayed using CellMinerCDB (https://discover.nci.nih.gov/SclcCellMinerCDB/) by plotting the same gene (expression, copy number or methylation), drug or microRNA on the x-Axis and the y-Axis. Expression of *Schlafen 11* (*SLFN11*), a gene whose expression is highly predictive of cytotoxic response to a broad range of DNA targeted agents including frontline treatments of SCLC (etoposide, topotecan, cis- and carboplatin) as well as drugs under investigation such as the poly(ADP-ribose polymerase) inhibitors (olaparib, niraparib, rucaparib, talazoparib) (Barretina et al., 2012; Farago et al., 2019; Gardner et al., 2017; Murai et al., 2019; Reinhold et al., 2017a; Zoppoli et al., 2012) measured by RNA-seq in the UTSW database shows a 0.92 Pearson’s correlation with its measured values by Affymetrix microarray in the NCI database (Figure 2B). *SLFN11* promoter DNA methylation measured by high resolution Illumina 850k arrays in the NCI database also shows a 0.9 Pearson’s correlation with its measured values by Illumina 450k microarray in the GDSC database (Figure 2C).

The other examples of cross-database analyses in Figure 2 are for *MYC*, which is commonly amplified and drives proliferation of a large fraction of SCLC (Dammert et al., 2019; Gazdar et al., 2017), *BCL2*, which encodes a canonical antiapoptotic protein targeted by Navitoclax (ABT-263) (Rudin et al., 2012) and for two SCLC drugs etoposide and topotecan. *MYC* amplification (by 850k methylome array in NCI) is correlated with its overexpression (by RNA-seq in CCLE) (Figure 2D). The activity of navitoclax is correlated with *BCL2* expression, suggesting BCL2 addiction for the cells overexpressing *BCL2*. Drug activity data for etoposide are correlated in the NCI and CTRP databases (note that drug activity was measured by different assays in each database; Rajapakse et al. (2018)). The cells most sensitive or resistant to etoposide overlap for topotecan.

Integrating the broader CellMinerCDB database (http://discover.nci.nih.gov/cellminercdb) of over 1000 cell lines including 74 and 53 SCLC cell lines in GDSC and CCLE (see Figure 1A) allows comparison between tissue of origin using CellMinerCDB (Rajapakse et al., 2018). For instance, the expression of *MYC* is correlated with the replication processivity factor *PCNA* (<u>P</u>roliferating <u>C</u>ell <u>N</u>uclear <u>A</u>ntigen) in SCLC (green) vs. other tissues including NSCLC (red), consistent with the replicative genotype of SCLC based on their high *PCNA* expression (alike leukemia and lymphoma cell lines) compared to NSCLC. Yet, high *MYC* expression is a feature of both the SCLC and NSCLC cell lines.

### SCLC Methylome

Two prior studies described the DNA methylation profiles of SCLC with limited data for established cell lines; 18 cell lines were examined by Kalari et al. (2013) and 7 by Poirier et al. (2015) together with primary tumors and PDX samples. Here we determined the methylome of the 66 cell lines of the NCI and processed the methylome data for the whole 985 GDSC cancer cell line dataset including its 61 SCLC cell lines. The data are highly reproducible in the two datasets for the 43 common cell lines (see Figure 2A and 2C). Thus, the SCLC-CellMiner resource provides the largest promoter methylation database for a total of 84 individual SCLC cell lines (43 common + 23 specific to NCI-SCLC + 18 specific to GDSC).

#### Globally low methylation levels of SCLC cell lines

Global methylation levels showed marked differences between the SCLC cell lines and the other cancer cell lines from different histologies. The median level of global methylation of the SCLC cell lines is the lowest compared with 21 subtypes of cancers (Figures 3A-B), which may reflect their intrinsic plasticity and stemness.

**Figure 3.**
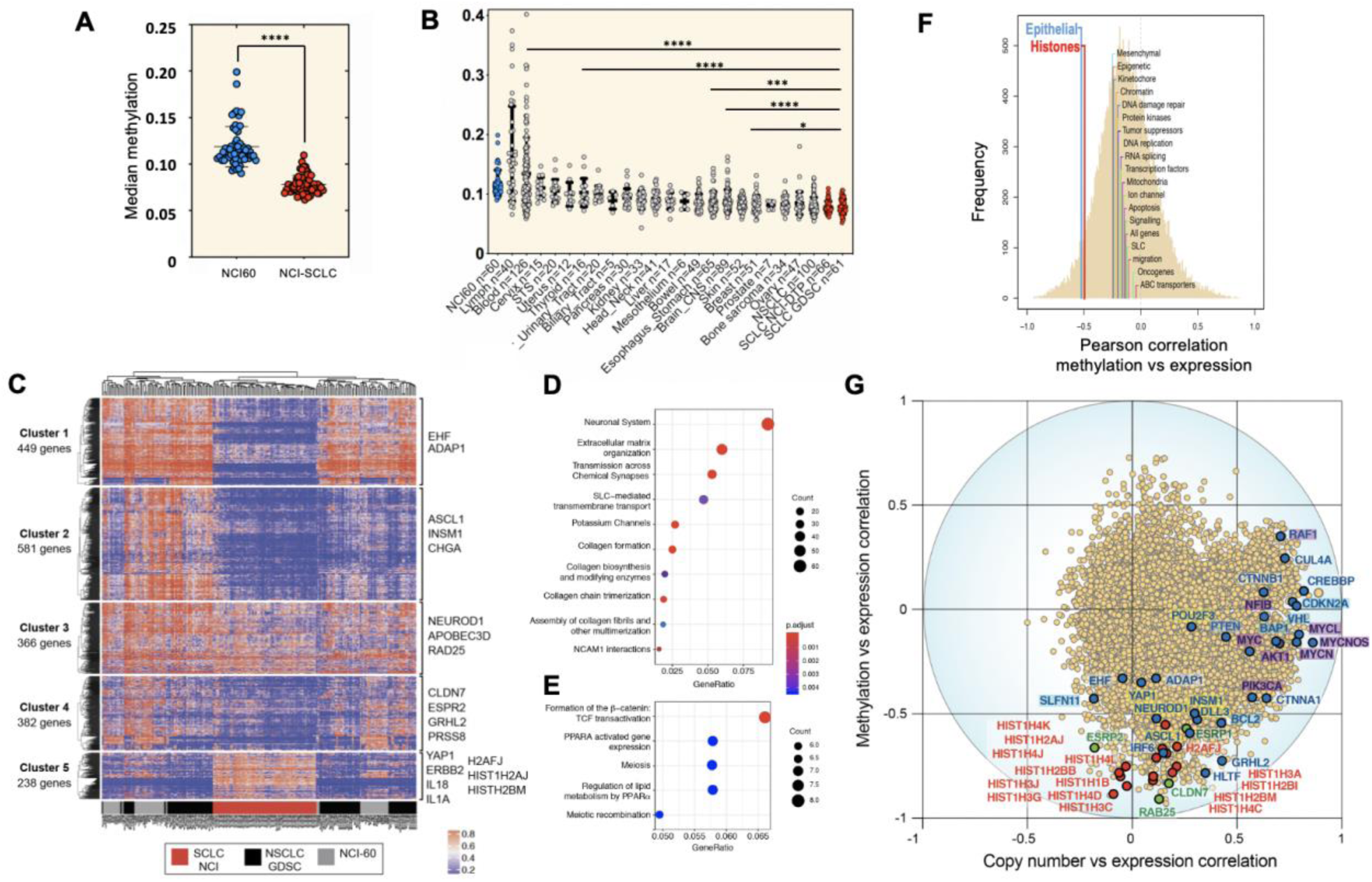
Methylation profile of SCLC cell lines. (**A**) SCLC cell lines exhibit low global methylation level compared to the non-SCLC of the NCI60 and of the GDSC (**B**). Each point represents the median methylation level of individual cell lines for the total set of 17,559 genes. Twenty one different cancer subtypes are ranked according their global methylation level. SCLC cell lines from two different sources (NCI and GDSC; in red) show the lowest global level of methylation. (**C**) Comparison of the methylation profiles between SCLC cell lines (red bar at bottom), NSCLC cell lines included in the GDSC and NCI-60 (black bar), and non-lung cancer cell lines from the NCI-60. The heatmap displays the median level of methylation of 2,016 genes with high dynamic range (genes with a standard deviation > 0.25 across the data sources) in the cell lines from SCLC-NCI (N=66), NSCLC-GDSC (N=75) and the NCI60 (N=60). Dark blue and dark red represent lowest and highest methylation median levels, respectively. Subtypes of the cell lines is indicated at the bottom (SCLC: red, NSCLC: black and NCI60: grey). SCLC cell lines represent one independent and distinct cluster. Among the 5 gene clusters, 3 show low methylation and one high methylation levels in SCLC. Examples of key SCLC genes are indicated at right. Details are provided in Supplemental Table S4. (**D**) Pathway analysis for clusters 1 & 2. (**E**) Pathway analysis for cluster 5. (**F**) Functional categories with significant correlation between gene transcript expression and DNA methylation. The figure shows histograms of the distribution of correlations of 17,144 transcript expression and DNA methylation data for the NCI-SCLC cell lines (N = 66). Median values are shown for the transcript expression versus DNA methylation level correlations of 20 functional groups of genes (defined in Supplementary Table S5). The x-axis are the Pearson correlations of the transcript expression versus the DNA methylation values, and the y-axis is the frequency. (**G**) Correlations between gene expression and predictive values of DNA copy number (X-axis) vs DNA methylation (Y-axis). An R value of 0 indicates no predictive power. R value of 1 or −1 and +1 indicate perfect negative and positive predictive power, respectively. Each point represents one of a total of 14,046 genes analyzed. Oncogenes and tumor suppressor genes (highlighted in purple and in blue, respectively) are primarily driven by copy number. Histone genes (red), and epithelial genes” (green) are primarily driven by DNA methylation (see Supplementary Table S5 for details. SCLC key genes (ASCL1, NEUROD1, POU2F3 and YAP1) are also indicated.

Yet, expression of some key SCLC genes is driven by promoter methylation, such as *ASCL1* and *NEUROD1* (Supplemental Figure S1). Cells not expressing those genes tend to be overmethylated. Conversely, cells expressing *ASCL1, NEUROD1, YAP1* and *POU2F3* have no significant promoter methylation. Yet, hypermethylation is not detetable in a number of cell lines that do not express those genes implying that further studies are warranted to examine other epigenetic markers (likely histone marks) as regulators of SCLC gene expression.

#### SCLC cell lines have a distinct methylome

To determine the methylation signature of the SCLC cell lines and differences with other cancer types, we compared the DNA methylation profiles of the NCI-SCLC to the methylation profiles of the NCI-60 (which includes 7 tissues of origin with 6 NSCLC cell lines but no SCLC cell lines) and of 75 NSCLC cell lines of the GDSC. After selecting a total of 2,016 genes with the most variable methylation (standard deviation > 0.25), we performed hierarchical clustering (Figure 3C). All the SCLC cell lines segregated together, while the NSCLC cell lines (N = 83 from GDSC and NCI-60) formed 4 clusters interrupted by SCLC cell lines (Figure 3C). The 2,016 genes clustered into three main groups: i) genes hypomethylated in SCLC cell lines (clusters 1,2 and 4), ii) genes hypermethylated in SCLC cell lines (cluster 5), and iii) genes with high methylation range in all cell lines independent of their tissue of origin (cluster 3). The detailed list of the genes in each cluster is provided in Supplemental Table S3.

Pathway analysis of the 1,030 specifically hypomethylated genes (clusters 1 + 2) shows an enrichment of neurological as well as extracellular matrix (ECM) pathways (Figure 3D and Supplemental Table 3), consistent with the neuroendocrine and cell aggregation features of the classic SCLC cell lines (Gazdar et al., 2017). Among these neuroendocrine (NE) genes, figure *ASCL1, CHGA* and *INSM1*, which is consistent with their expression. Many genes involved in epithelial–mesenchymal transition (EMT) (Kohn et al., 2014) also tend to be hypomethylated in the SCLC cell lines including *CLDN7, ESRP2, MARVELD2, PRSS8, ST14, IRF6, GRHL2, CLDN4, EHF, ADAP1* and *CMTM3*. Most of the EMT genes belong to cluster 4 and are also hypomethylated in the NSCLC cell lines.

Analysis of the 238 genes selectively hypermethylated in SCLC (cluster 5) shows a significant representation of the beta-catenin/Tcf transaction and Wnt signaling pathway as well as genes involved in lipid metabolism by peroxisome proliferation-activated receptor alpha (PPARα) (Figure 3E). *YAP1* and *ERBB2* are also hypermethylated in most cell lines, as well as a large fraction of the canonical histone genes.

#### Expression of histone and epithelial genes is highly driven by methylation in SCLC cell lines

To further determine gene categories driven by promoter methylation, we compared the gene expression and methylation pattern of functional groups (Reinhold et al. (2017c); Supplemental Table S4). Two functional gene categories showed strong correlation between methylation and expression: epithelial and histone genes (Figure 3F), with 25 and 75 genes, respectively. The median correlation was - 0.53 for the epithelial genes and - 0.50 for the histone genes.

Analysis of individual genes (Figure 3G) confirmed that histone genes are dominantly regulated by methylation in SCLC. Among the 62 canonical histone genes with available data, 21 belong to H2A core histone family, 18 to H2B core histone family, 14 to H3 core histone family, 13 to H4 core histone family and 9 to the H1 linker family. Among the 13 non-canonical histones, 4 are replication independent histones (*H1F0, H1FNT, H1FOO, H1FX*) and replacements of H1 histone. Their transcription is independent of DNA replication and they are expressed throughout the cell cycle in a tissue specific manner. The remaining are variants from core histones (*H2AFJ, H2AFX, H2AFY2, H2AFY, H3F3C, H3F3B, H2AFV, H2AFZ*). Unlike canonical histones that function primarily in genome packaging and gene regulation, variant histones distinct function including DNA repair, meiotic recombination and chromosome segregation (Buschbeck and Hake, 2017). Canonical histones showed the highest correlation between expression and methylation suggesting that epigenetic regulation of canonical histone is a feature of SCLC carcinogenesis. On the contrary, we find that the expression of the non-canonical histones is inconsistently driven by methylation suggesting a higher dynamic state across the SCLC cell lines.

Detailed analysis of the macroH21 variant *H2AFY* using the RNA-seq data from the UTSW database revealed that SCLC cell lines predominantly express the macroH2A1.2 variant compared to the macroH2A1.1 variant. The macroH2A1.2 splice variant is known to promote homologous recombination and is essential for proliferation (Kim et al., 2018). This finding is consistent with the characteristically high proliferation of SCLC cell lines, which is regulated by methylation and epigenetics in addition to *RB1* and *TP53* inactivation and *MYC* oncogene overexpression.

### SCLC DNA Copy Number vs Methylome as Drivers of Gene Expression

To determine how gene copy number and promoter methylation account for gene expression in the SCLC cell lines, we analyzed whole-genome DNA copy number data and correlated the expression of each gene with DNA copy number (x-axis) and methylation (y-axis) (Figure 3G) (Reinhold et al., 2017c). 84% of the genes showed positive correlation with copy number and 65% negative correlation with DNA methylation. Consistent with the pathway analyses (Figure 3F), epithelial (green) and histone genes (red) were most consistently driven by promoter methylation. Correlations for individual genes between methylation and expression can be readily checked using SCLC-CellMiner (https://discover.nci.nih.gov/SclcCellMinerCDB/). Snapshot examples of genes driven by methylation (*NEUROD1, ASCL1, POU2F3, YAP1, SLFN11, SMARCA1, SOX1* and *CGAS*) are presented in Supplemental Figure S1. Genes exhibiting low or no expression did not show a consistent correlation with promoter hypermethylation, consistent with diverse mechanisms for inhibiting gene expression. For each gene, CellMinerCDB allows the identification of cell lines with methylation-dependent and independent gene expression for further molecular and mechanistic studies.

Unlike the histone and epithelial genes, which are primarily driven by DNA methylation, the expression of key SCLC growth-driving genes, such as the oncogenes (*MYC, MYCL, MYCN, AKT1*) the tumor suppressor genes (*CDKN2A, BAP1, VHL*) and the chromatin remodeler genes *(EP300* and *CREBBP*) are mainly driven by DNA copy-number alterations (Figure 3G). R values for any gene of interest (with data) are provided in Supplemental Table S5. Examples of CellMinerCDB snapshots are provided in Supplemental Figure S2 for *MYC, MYCL* and *MYCN, BAP1* and *VHL*, whose expression is driven by copy number changes but not by DNA methylation.

### SCLC-Global Integrates the Transcriptome of all 116 SCLC Cell Lines

To take advantage of all 116 cell lines with expression data by microarray or/and RNA-seq across the five data sources (Figure 1), we regrouped them by normalization using Z-score to remove dataset batch effects. Principal component and correlation analyses validated the approach (Supplemental Figure S3A-C). The data are available under “*SCLC Global*” at https://discover.nci.nih.gov/SclcCellMinerCDB/ in the pull-down tab for the “x- and y-Axis Cell Line Set”. For instance, the correlation for ASCL1 expression in the “SCLC-Global” vs SCLC NCI/DTP gives a Pearson’s correlation coefficient of 0.99 with a p-value=1.9e-55. SCLC-Global offers many other features including cross-correlation with other databases for DNA methylation, DNA copy number, DNA Mutation, MicroRNA or Drug Activity.

*SCLC-Global* gene expression tools can be used to retrieve all the genes correlated with the expression of any given gene. For instance, for the *MYCN* gene (Supplemental Figure S4A-C), the top correlate (Pearson’s correlation coefficient 0.967) is *MYCNOS*, the *MYCN Opposite Strand* antisense RNA. The data for individual cell lines can also be visualized by plotting *MYCNOS* against *MYCN* in the SCLC-Global database (Supplemental Figure S4B). Notably plotting *MYCN* vs *MYCNOS* in the CCLE database using CellMinerCDB extends the finding that MYCN is co-expressed with its antisense RNA in both SCLC and brain tumor cell lines (Pearson’s correlation coefficient 0.81; Supplemental Figure S4C).

### SCLC Molecular Signatures: NE, NAPY and MYC Signatures

Next, we tested the *SCLC-global* gene expression data to explore and validate the recently established molecular signatures of SCLCs (Rudin et al., 2019). As indicated previously, SCLC can be classified as neuroendocrine (NE) or non-neuroendocrine (non-NE) with only 10-25% being non-neuroendocrine as defined by lack of expression of key neuroendocrine markers (Gazdar et al., 2017; Gazdar et al., 1985; McColl et al., 2017; Rudin et al., 2019; Zhang et al., 2018).

Using the *SCLC-Global* dataset, we scored the 116 cell lines based on the classification of Gazdar and coworkers (Augustyn et al., 2014; Zhang et al., 2018), which uses the expression values of 50 genes to calculate a NE score. This NE score is highly correlated with the expression of *SYP* (encoding Synaptophysin), *CHGA* (encoding Chromogranin A), and *INSM1* (encoding Insulinoma Transcriptional Repressor) (Figure 4A), which are used in routine diagnosis to establish the NE characteristics of SCLC biopsies. To explore the selectivity of these genes for SCLC cell lines, we examined the large collection of cell lines of the GDSC and CCLE (Rajapakse et al., 2018). *CHGA, INSM1* and *SYP* were selectively expressed both in SCLC and brain tumors, which is consistent with the neuronal differentiation of SCLC (Supplemental Figure S5A-B). Moreover, the NE-SCLC cell lines, which can be readily labeled in SCLC-CellMinerCDB under the “Select Tissues to Color” tab, have significantly higher levels of expression of *CHGA* and *SYP* compared to non-NE cell lines (Supplemental Figure S5C).

**Figure 4.**
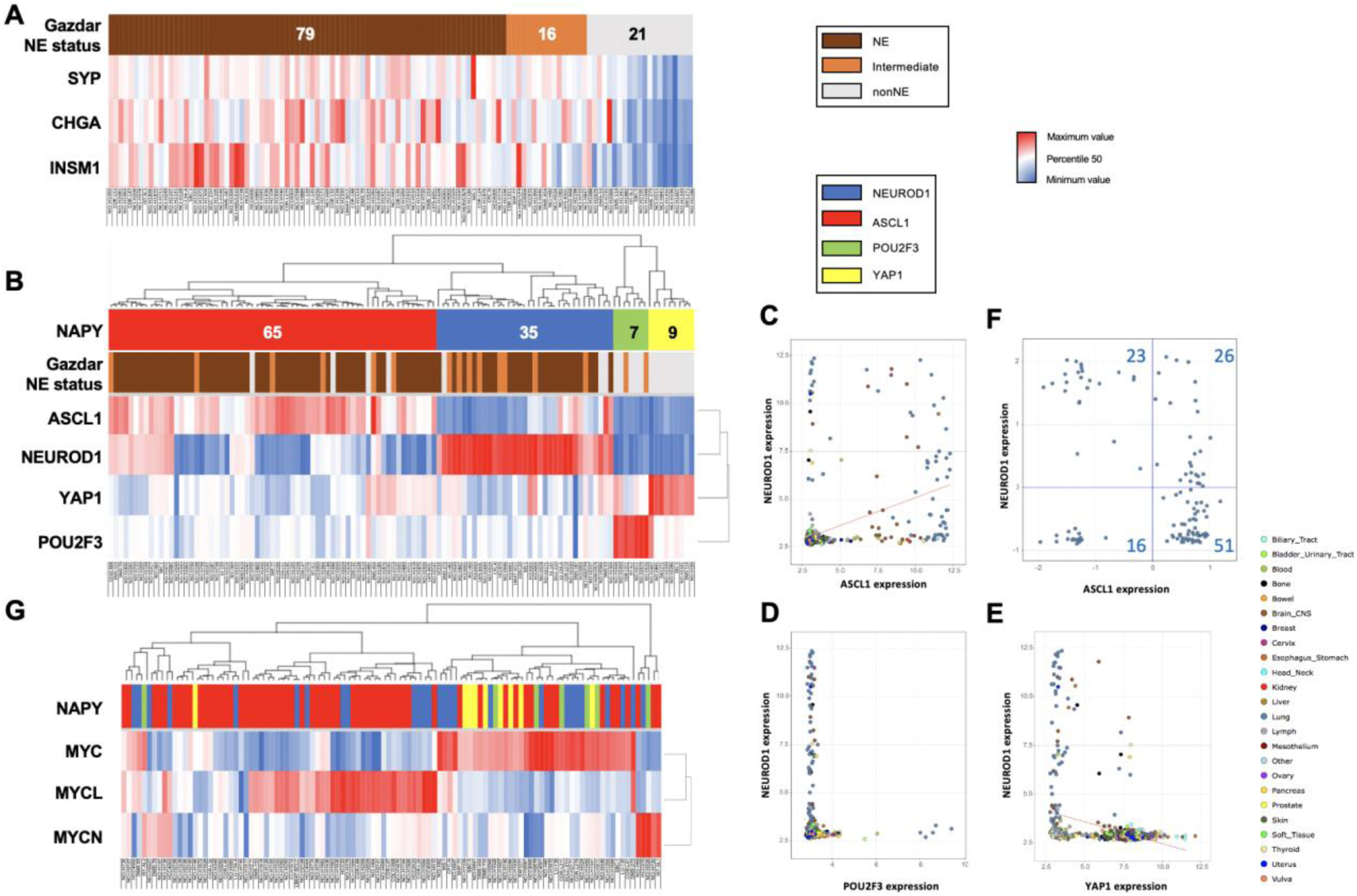
SCLC genomic molecular classifications. (**A)** Neuroendocrine *versus* non-neuroendocrine classification based on the expression of 50 genes (Gazdar et al., 2017). Neuroendocrine (NE; in dark brown) and non-neuroendocrine status (nonNE; in grey) scores are represented for each cell line (N = 116). In light brown are the cell lines with an intermediate score. Numbers at the top correspond to the number of cell lines in each group. Expression of the clinical histological biomarkers *CHGA, SYP* and *INSM1* is included. They were obtained after normalization by Z-score (see Supplemental Figure S2). Red and blue correspond to high and low gene expression, respectively. Detail are provided Supplementary Table S3. (**B**) Classification based on *NEUROD1, ASCL1, POU2F3* and *YAP1* (NAPY) expression (Rudin et al., 2019). The heatmap displays expression of the NAPY genes in the overall 116 SCLC cell lines of SCLC-CellMiner. Expression values across the 5 data sources were obtained after normalization by Z-score (see Supplemental Figure S2). Complete distance hierarchical clustering shows the expected 4 groups of cell lines. *ASCL1* (N = 65) and *NEUROD1* (N = 35) high-expressor cell lines are considered as NE-SCLC cell lines and *POU2F3* (N = 7) and *YAP1* (N = 9) cell lines, non-NE-SCLC cell lines. The Gazdar classification is included for comparison. Details are provided in Supplementary Table S3. (**C**) *NEUROD1* and *ASCL1* are specific for both SCLC and brain tumor cell lines. Expression of *ASCL1* versus *NEUROD1* in the GDSC database and processed with CellminerCDB. Each point represents a cell line (N = 986). (**D**) Common co-expression of *NEUROD1* (y-axis) and *ASCL1* (x-axis) in the 11 SCLC. Each point represents a cell line. (**F**) *POU2F3* is selectively expressed in SCLC but not in brain tumor cell lines (N=986 from GDSC processed with CellMinerCDB). (**G**) *YAP1* expression is not specific to SCLC. *YAP1* exhibits a high range of expression across the different subtypes of cancer cell lines of the GDSC database (N=986). Plots in panels E-F are snapshots from CellMinerCDB (http://discover.nci.nih.gov/cellminercdb). (**G**) Classification based on MYC genes expression. The heatmap displays expression of *MYC, MYCL* and *MYCN* in 106 SCLC cell lines across the 5 data sources after normalization by Z-score (see Supplemental Figure S2). The figure also provides the NAPY classification for each cell lines. Details are in Supplementary Table S4.

Rudin et al. (2019) proposed a more detailed molecular classification based on the expression of four transcription factor genes: *NEUROD1* and *ASCL1* for neuroendocrine, and *YAP1* and *POU2F*3 for non-neuroendocrine SCLCs (Figure 4B, Supplemental Table S6). Compared to the other cancer cell lines in the GDSC-CellMiner database, the highest expression of *NEUROD1* and *ASCL1* is found in SCLC and brain tumors (Figure 4C), while *POU2F3* expression is rare and limited to SCLC cell lines (Figure 4D). In contrast, *YAP1* is not limited to SCLC and is expressed in a wide range of cancer types (except blood and lymphoid tumors) in addition to the non-neuroendocrine SCLC (Figure 4E). Differential expression of the 4 transcription factors (“NAPY” classification for short) across the SCLC-Global database of 116 cell lines clearly distinguishes the four subtypes of SCLC cell lines (Figure 4B), with similar proportions as reported by Rudin et al. (2019) across tumors and cell lines. *ASCL1* expression is commonly associated with *NEUROD1* expression (Figure 4B), indicating that a significant fraction of NE-SCLC cells have dual expression of *ASCL1* and *NEUROD1*. Figure 4F shows that 63% of the *ASCL1*-expressing cells co-express *NEUROD1* and 47% of the *NEUROD1*-expressing cells co-express *ASCL1*.

The NE and NAPY classifications show high concurrence across the *SCLC-Global* cell lines (93.9% agreement with Cohen’s kappa of 0.79 after excluding intermediates; Figure 4) with the three NE genes *CHGA, SYP* and *INSM1* most significantly overexpressed in the NEUROD1 and ASCL1 subgroups compared to the POU2F3 and YAP1 subgroups of non-NE SCLC cell lines (Supplemental Figure S5D-E).

The three MYC-genes *MYC, MYCL* and *MYCN* play key roles in SCLC carcinogenesis. *MYCL* was discovered as being selective amplified in SCLC (Johnson et al., 1987; Nau et al., 1985). Close to 80% of the SCLC cell lines highly express one of the three MYC genes with *MYC* and *MYCL* being the most prevalent (Figure 4G). Notably, and as noted previously, cells overexpressing one of the MYC-genes are negative for the two other MYC genes, indicating a mutually the mutually exclusive expression of the 3 MYC genes. Also, the non-NE SCLC cell lines (SCLC-Y and SCLC-P) express low *MYCL* and *MYCN* compared to the NE-SCLC (SCLC-A and SCLC-N) and YAP1 cells, which selectively express *MYC* but neither *MYCL* nor *MYCN* (Figure 4G and Supplemental Figure S6A-B).

### SCLC Transcriptional Networks Focusing on the ASCL1, YAP/TAZ and NOTCH Pathways

Because the four NAPY genes (*NEUROD1, ASCL1, POU2F*3 and *YAP1*) are transcription factors, we performed transcription network analyses (Kohn et al., 2006) in connection with the NAPY classification. Snapshots are presented in Supplemental Figure S4A-B, S5C,S6C and S7A&C for the “Univariate Analyses” and in Figure 5B & D and Supplemental Figure S5E for “Multivariate Analyses” (https://discover.nci.nih.gov/SclcCellMinerCDB/).

**Figure 5:**
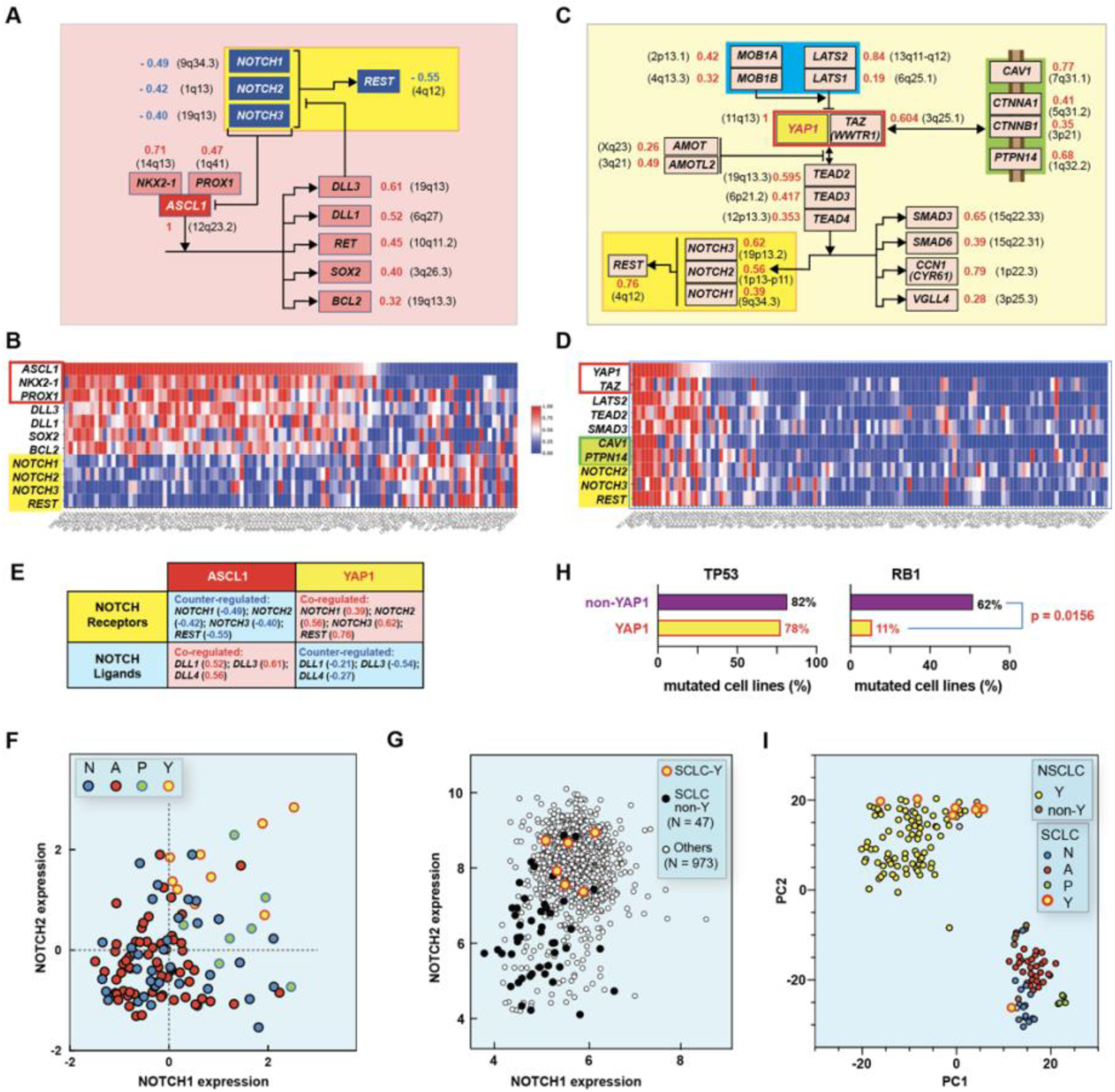
Integration of the transcriptional networks of the neuroendocrine ASCL1 and non-neuroendocrine YAP1 SCLC cell lines with the NOTCH pathway. (**A**-**D**). ASCL1 (panels A-B) and YAP1 (panels C-D) networks. Panel A shows the highly significant correlations between *ASCL1* expression and its molecular transcriptional coactivators *NKX2-1* and *PROX1*, and some of its downstream transcriptional targets (bayonet arrows). Numbers to the right indicate the significantly positive Pearson’s correlations coefficients (red) and chromosome locations (black in parenthesis) obtained from Miner Global (https://discover.nci.nih.gov/SclcCellMinerCDB). The NOTCH receptor network (blue boxes) with its transcriptional target *REST* are shown at the top of the panel (yellow box). Negatively significant Pearson’s correlations coefficients (blue) and chromosome locations (black in parenthesis) obtained from SCLCcellMiner Global (https://discover.nci.nih.gov/SclcCellMinerCDB) Panel B: visualization of the correlations between *ASCL1* expression and the indicated genes corresponding to those shown in panel A. Note the counter-expression of the NOTCH receptor pathway (yellow highlight) with respect to *ASCL1* expression. The image is a snapshot obtained using the multivariate analysis tool of SCLCcellMiner using the Global dataset of the 116 cell lines. Panels C and D. Same as panels A and B except for *YAP1* across the 116 SCLC cell lines of SCLCcellMiner. Note the positive correlation between *YAP1* expression and the NOTCH receptor pathway (see text for details). (**E**) Negative correlations between the NOTCH receptors and ligands and *ASCL1* vs *YAP1* across the 116 cell lines of SCLCcellMiner. Pearson’s correlation coefficients with respect to *ASCL1* (2nd column) and *YAP1* (3rd column) are indicated in parenthesis. They can be obtained using the Global dataset of the 116 cell lines of SCLCcellMiner. (**F**) Correlation between *NOTCH1* and *NOTCH2* across the Global dataset of the 116 cell lines of SCLCcellMiner. *YAP1* cells show significantly highest expression of both *NOTCH1* and *NOTCH2*. (**G**). Correlation between *NOTCH1* and *NOTCH2* across the 1036 cell lines of the CCLE. The SCLC-YAP1 have highest NOTCH (see inset for annotations). (**H**) SCLC-YAP1 cells have significantly reduced frequency of RB1 mutations. Only one SCLC-YAP1 cell line (NCI-H196) shows RB1 mutation whereas 7 of the 9 SCLC-YAP1 show TP53 mutations. Data were compiled from the 116 cell lines of SCLC-CellMiner Global (**I**). tSNE clustering plot using gene expression data of 60 SCLC and 100 NSCLC cell lines (microarray; GDSC data source). Each dot represents a sample and each color represents the type of the sample (see inset).

Figure 5A summarizes our analyses of the ASCL1-NOTCH genomic transcriptional network based on our molecular interaction map (MIM) conventions (Kohn et al., 2006) (https://discover.nci.nih.gov/mim/index.jsp). As a pioneer transcription factor, ASCL1 binds E-box motifs (as NEUROD1) to promote chromatin opening and the activation of neuronal genes. Notably both *NKX2*.1 and *PROX1*, whose encoded polypeptides function together as transcription cofactors with ASCL1 are highly significantly co-expressed with *ASCL1* in the SCLC cell lines, and this co-expression is not due to the presence of those genes on the same chromosomes (Figure 5A), indicating upstream regulatory transcriptional control with the likely implication of super-enhancers. As expected, the transcriptional targets of ASCL1 were co-expressed with *ASCL1* (Figure 5A-B). One of those known targets, *BCL2* is positively correlated not only with *ASCL1* expression (Figure 5A-B) but also with *POU2F3*, whereas *BCL2* expression was found negatively correlated with *NEUROD1* expression (Supplemental Figure 7A-B). Expression of the cancer-driving genes *RET, SOX1, SOX2, FOXA1* and *FOXA2* are also highly correlated with *ASCL1* expression (Figure 5A-B).

*DLL3*, another established transcriptional target of ASCL1 and a known inhibitor of the NOTCH pathway was found highly significantly correlated with *ASCL1* (r = 0.61; p = 4.05e-13; Figure 5A). Analysis of the NOTCH pathway whose inactivation is crucial in NE-SCLC (Gazdar et al., 2017; Leonetti et al., 2019; Ouadah et al., 2019) using the *SCLC-Global* database showed that the 3 NOTCH transcripts (*NOTCH1, NOTCH2* and *NOTCH3*) are jointly downregulated in the *ASCL1* SCLC cell lines (Figure 5A-B). Functional downregulation of the NOTCH pathway is consistent with the highly significantly negative correlation (r = -0.545; p = 2.45e-10) between *ASCL1* and *REST*, the transcriptional target of NOTCH (Figure 5A). Notably, the NEUROD1 subset of NE-SCLC (SCLC-N) did not show a significant correlation between *NEUROD1* and *DLL3* expression (r = -0.18; NS) (Supplementary Figure S7C-D), providing no evidence that *DLL3* overexpression acts to down-regulate the NOTCH pathway in SCLC-N cell lines. Hence, in the SCLC-A cell lines, the negative correlation between *ASCL1* and NOTCH genes could be related to the direct transcriptional inactivation of *ASCL1* by NOTCH3 (Figure 5A).

Of the 116 SCLC cell lines in SCLC-CellMiner, nine belong to the YAP subset (see Figure 4B&E). Because expression of YAP (*YAP1*) is also a feature in a wide variety of solid tumor cells (see Figure 4E), and YAP and its regulatory Hippo signaling pathway are the focus of many ongoing studies, we explored the YAP transcriptional network in the SCLC cell lines (Figure 5C). The first notable finding is that *YAP1* expression is highly correlated with the expression of its heterodimeric partner *TAZ* (encoded by the *WWTR1*/*TAZ* gene) both in the *SCLC-Global* dataset (Figure 5C-D) and across the 986 cell lines of the GDSC (Supplementary Figure S8). This finding suggests a master transcriptional regulator upstream of both genes or *YAP1* acting as super-enhancer, as both genes are on different chromosomes (Figure 5C; chromosome location indicated in italic and parenthesis).

Next, we explored the Hippo pathway, which acts as a negative regulator of YAP/TAZ and is commonly inactivated in solid tumors (Dasgupta and McCollum, 2019; Ma et al., 2019; Totaro et al., 2018). Expression of both *LATS2* and *LATS1*, which encode the core kinase of the Hippo pathway and negatively regulate YAP by sequestering phosphorylated YAP in the cytoplasm, are significantly positively correlated with *YAP1* expression (Figure 5C-D). This unexpected finding suggesting a negative feedback loop is additionally supported by the fact that the transcripts of *MOB1A* and *MOB1B*, the cofactors of LATS1/2, are also positively correlated with *YAP1* (Figure 5C-D). Moreover, the transcripts of the negative regulators of YAP, *AMOT* and *AMOTL2*, which are released by depolymerized F-actin and sequester YAP from its nuclear translocation, are also significantly positively coregulated with *YAP1* (Figure 5C-D) (Dasgupta and McCollum, 2019; Wang et al., 2019). Together, these results demonstrate that the YAP-SCLC cell lines co-express both YAP/TAZ and its negative regulator genes driving the Hippo pathway, and suggest an equilibrium (“metastable”) state where the Hippo pathway remains active to potentially negatively regulate YAP/TAZ in the Y-SCLC cells.

YAP/TAZ functions as a direct activator of the TEAD transcription factors (encoded by *TEAD2/TEAD3/TEAD4*), whose expressions are highly significantly coregulated with YAP1 (Figure 5C). As expected, the transcriptional targets of the TEADs are also significantly correlated with *YAP1* expression, some of which are included in Figure 5C (bottom section. Others can readily be found and discovered using the “Compare Pattern” of SCLC-CellMiner using the “Compare Pattern” of SCLC-CellMiner with TEAD or YAP1 as “seeds”. Among those are the cancer- and growth-related *SMAD3* and *SMAD5* genes, *CCN1*/*CYR61*, which encodes a growth factor interacting with integrins and heparan sulfate, and *VGLL4* (Figure 5C, bottom right and Figure 5D).

The NOTCH pathway is also a known transcriptional target of YAP/TAZ and the TEADs (Totaro et al., 2018). Consistent with this, we found a high positive correlation between *YAP1* the NOTCH receptor transcripts *NOTCH1, NOTCH2, NOTCH3* as well as the NOTCH transcriptional target *REST*, demonstrating the functional activation of the NOTCH pathway in SCLC-Y cells (Figure 5C-E). By contrast, and consistent with the biology of the NOTCH pathway, 4 of the 5 NOTCH ligands, *DLL1, DLL3, DLL4* and *JAG2*, which act as negative regulators of the NOTCH receptors (Andersson et al., 2011) are significantly negatively correlated with *YAP1* (Figure 5E). The results of these analyses support the conclusion that the NOTCH pathway is “on” in the SCLC-Y cells. By contrast, in the SCLC-A cells, the opposite is observed: the transcripts for the NOTCH receptors and the NOTCH ligands are negatively and positively correlated with the expression of *ASCL1* (Figure 5E and Supplementary Figures S9A). Notably, the SCLC-P cells also show a positive correlation between the NOTCH receptor and *REST* effector transcripts and *POU2F3* (Figure 5F and Supplementary Figure S9A and S10A). These analyses demonstrate a clear difference between the NE-SCLC (SCLC-N & -A) and the non-NE-SCLC (SCLC-P & -Y) with respect to the NOTCH pathway; with the pathway “off” in the NE subset (N & A) and “on” in the non-NE subset (P & Y).

Global analyses of the NOTCH pathway across the 1,036 cell lines from 22 different tissue types of the Broad-CCLE collection (Figure 5G and Supplementary Figure S9B-C) show that *NOTCH2* and *NOTCH3* expression are coregulated in many tumor types, especially brain, lung, lymph, thyroid, pancreas and uterus (Supplementary Figure S9B-C) and that the NE-SCLC cell lines are characterized by lowest NOTCH expression (Figure 5G and Supplementary Figure S9B). By contrast, the SCLC-Y- and -P cells are found among the NOTCH expressing cells. Of note, analyses of the NOTCH pathway activity measured by REST expression shows that the SCLC-Y cells cluster with the NSCLC cell lines (Figure 5G and Supplementary Figure S10B).

### Transcriptome of SCLC-Y Cells is Common with NSCLCs and Specific to this Subgroup

To further examine the relationship between the SCLC-Y cell lines and the NSCLC cell line, we performed principal component and other dimension reduction analyses with respect to the whole transcriptome data (Figure 5I). tSNE (t-distributed Stochastic Neighbor Embedding) is a method to highlight strong patterns in a dataset by reducing the dimensionality of a dataset while preserving as much ‘variability’ as possible. We performed tSNE analysis using gene expression data between NSCLC (N = 100) and SCLC (N = 60) cell lines from the GDSC data source to identify clusters of subgroups. This approach segregated the SCLC-Y together with the NSCLC cell lines. The other SCLC cell lines (SCLC-A, SCLC-N and SCLC-P) formed a distinct cluster. Also, among the few NSCL cancer cell lines clustering with the NAP-SCLC were carcinoids of the lung and one misannotated cell line. These data support that SCLC-Y cell lines are a distinct entity among the SCLC subtypes and potentialy related to NSCLC.

Another characteristic of the SCLC-Y cell lines is the significantly low *RB1* mutations (only one cell line among 9 showing *RB1* mutation; Figure 5H). The SCLC-Y cell lines also showed significantly reduced activity of the replication transcriptional network with highest *RB1* expression and lowest *PCNA, MCM2* and *RNASEH2A* expression (Supplementary Figure S11A & D-F). Additionally, the SCLC-Y cells express the mesenchymal marker *VIM* as well as the cytoskeleton component and regulators *CNN2* (actomyosin and F-actin component) and the *AMOT* genes, which regulate cell migration and actin stress fiber assembly (Figure 5C, left and right) (Dasgupta and McCollum, 2019).

### Global Drug Activity Profiling Suggests Transcription Elongation Pathways as General Drug Response Determinant and Hypersensitivity of the SCLC-P Cell Lines

To explore potential connections between the NAPY classification and drug responses, we analyzed the drug sensitivity profiles of the 66 SCLC-NCI cell lines using 134 compounds with the highest activity range (> 0.09) (Polley et al., 2016). Unsupervised hierarchical clustering generated two groups of cell lines: those globally resistant to all drugs and those globally drug-sensitive, with a bimodal distribution (Figure 6A). No obvious relationship was observed for the neuroendocrine cell lines (SCLC-N and SCLC-A), which were distributed in both clusters. Yet, all three SCLC-P cell lines clustered together among the most globally drug-sensitive whereas the SCLC-Y cell lines tended to be among the most resistant cell lines.

**Figure 6:**
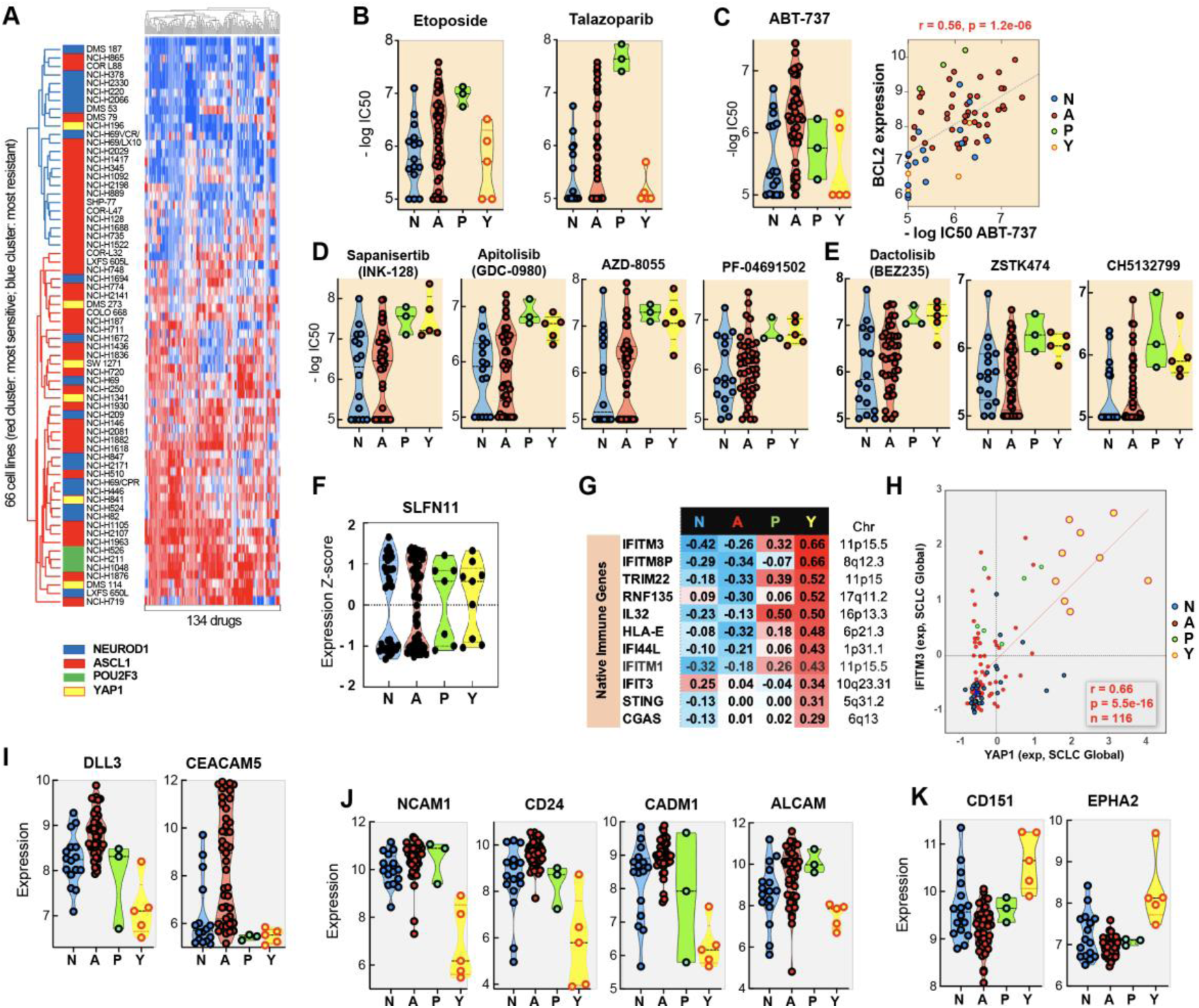
Therapeutic predictive genomic biomarkers for SCLC based on cancer cell lines drug response, gene expression and molecular NAPY classification. (**A**). Cluster image map showing the global response of the NCI-SCLC cell lines (N = 66) across 134 different drugs from a broad range of chemical classes and targets. Cell lines are listed in the middle column and their NAPY classification to the left. (**B**). POU2F3 cells are the most sensitive to etoposide and talazoparib while the YAP1 cell lines are the most resistant. (**C**). Selective activity of the BCL2-BCL-XL inhibitor in a subset of the ASCL1-SCLC cell lines (left) and highly significant correlation with BCL2 expression (right). (**D**). Selective activity of the mTOR/AKT inhibitors in a subset of the non-NE SCLC cell lines (POU2F3 = P; YAP1 = Y). (**E**). Selective activity of the PI3K inhibitors in the non-NE SCLC cell lines. (**F**). *SLFN11* expression across the 116 SCLC cell lines exhibits bimodal distribution in all 4 subtypes of SCLC and is a predictive biomarker for DNA damaging chemotherapeutic agents (https://discover.nci.nih.gov/SclcCellMinerCDB)] (see Supplemental Figure S12). (**G**). Selective expression of native immune pathway genes in the YAP1 SCLC (correlations between each of the NAPY genes and the listed native immune response genes are listed with colors reflecting significantly positive and negative correlations (red and blue, respectively). (**H**). Snapshot from SCLC-CellMiner illustrating the correlation between YAP1 and IFITM3 transcripts across the 116 cell lines of SCLC-CellMiner Global (see Supplemental Figure S13). (**I**). Selective expression of the DLL3 and CEACAM5 surface markers targeted by Rovalpituzimab tesirine (Rova-T) and Labetuzumab govitecan (IMMU-130), respectively, in the NE-SCLC cell lines (A preferentially) (see Supplemental Figure S13). (**J**). Potential surface biomarker targets for NE-SCLC and POU2F3 SCLC cells (N & A). (**K**). Potential surface biomarkers for non-NE YAP1-SCLC cells. Data in panels A-E and I-K are from the 66 cell lines from the NCI-DTP drug and genomic database.

Differential gene expression followed by enrichment pathway analyses was performed to determine potential differences between the most and least drug sensitive cell lines. The most significantly enriched pathway was the ribosomal and EIF2 signaling pathway, which was selectively activated in the sensitive compared to non-sensitive cell lines. EIF2 (Eukaryotic Translation Initiation Factor 2A) catalyzes the first regulated step of protein synthesis initiation, promoting the binding of the initiator tRNA to 40S ribosomal subunits. EIF2 factors are also downstream effectors of the PI3K-AKT-mTOR and RAS-RAF-MAPK pathways. The details of the analysis are provided in Supplemental Figure S12A-B. These results suggest that global drug response in SCLC is associated with active protein synthesis.

### Drug Activity Profiling in Relationship with the NAPY Classification

Both the ASCL1 (A) and NEUROD1 (N) subgroups showed a broad range of response to etoposide, topotecan and cisplatin, as well as to the potent PARP inhibitor talazoparib (Figure 6B and Supplemental Figure S12C). The most significant genomic predictor of response for these neuroendocrine SCLC-N & -A subgroup was *SLFN11* expression (Supplemental Figure S12C; https://discover.nci.nih.gov/SclcCellMinerCDB/), which is consistent with analyses performed across other tissue types (Barretina et al., 2012; Rajapakse et al., 2018; Zoppoli et al., 2012). The potential value of *SLFN11* expression as a predictive biomarker is also borne out by its highly dynamic and bimodal expression pattern (Figure 6F). Approximately 40% of the 116 SCLC cell lines of *SCLC-global* do not express *SLFN11* (Supplemental Figure S12D).

The SCLC-Y cell lines showed the greatest resistance to the standard of care drugs (etoposide, cisplatin and topotecan) (Figure 6B). This result is not limited to SCLC, as a highly significant drug resistance phenotype was observed between *YAP1* expression and response to etoposide and camptothecin across the database of the CCLE-CTRIP, which spans across a broad range of tissues of origin (Supplemental Figure S12E).

In addition to SLFN11, a predictive genomic biomarker of drug response is methylguanine methyltransferase (MGMT) for temozolomide (TMZ), which acts as a DNA methylating agent generating N7- and O6-methylguanines. MGMT removes O6-methylguanine, the most cytotoxic lesion. Cancer cells (typically glioblastomas) with MGMT inactivation are selectively sensitive to TMZ (Thomas et al., 2017). Analyses of the SCLC cell lines revealed lack of *MGMT* expression in 33% (N = 38) of the SCLC cell lines (Supplemental Figure S12D). Notably, the non-NE cell lines all expressed MGMT, indicating that the SCLC-P- and -Y cancer cells are predicted to be poor candidates to TMZ-based therapies (Farago et al., 2019).

To determine whether the NAPY classification predicts sensitivity to drugs not commonly used as standard of care for SCLC, we performed correlation analyses to identify the drugs that were significantly linked to a subtype among the 526 NCI compounds (Polley et al., 2016). The list of all the statistically significant drugs (p-value < 0.05; Kruskal Willis test) is provided in Supplemental Table S7). Eighteen drugs were highly subtype-specific (p-value < 0.01; Kruskal Willis test). Among them, 7 are PI3K-AKT-mTOR inhibitors and all of them show a higher activity in the non-NE cell lines (SCLC-Y and SCLC-P) (Figure 12D-E). The SCLC-P and -Y cell lines are also more sensitive to multi-kinase inhibitors including dasatinib or ponatinib. One agent was found specifically active in *ASCL1* high expressing cell lines: ABT-737, a *BCL2* inhibitor (Figure 6C). Analyzing the GDSC, CCLE and CTRP (https://discover.nci.nih.gov/SclcCellMinerCDB/) showed that all BCL-2 inhibitors are most efficient in the SCLC-A cell lines, while the SCLC-Y cell lines are consistently resistant. The high sensitivity of the SCLC-A cell lines is consistent with the highly significant correlation between BCL2 expression and the activity of ABT-737.

### Immune Pathways are selectively expressed in the YAP1 Subgroup of SCLCs

Although immune checkpoints inhibitors (ICI) have been approved in SCLC, the benefit in an unselected patient population is modest with approximately 2-month improvement in median overall survival when immunotherapy was added to first-line platinum and etoposide.

To explore the activity of the immune pathways in the 116 cell lines of *SCLC-Global* and the potential value of the NAPY classification for selecting SCLC patients likely to respond to immune checkpoint inhibitors, we explored the transcriptome of the cell lines by focusing on a subset of established native immune response and antigen-presenting genes. Figure 6G-H shows the unique characteristics of the SCLC-Y cell lines. Indeed, they are the only subset expressing innate immune response genes and for which expression of those genes such as the innate immune effector genes *CGAS* and *STING*, the antigen-presenting HLA gene (*HLA-E*) and the interferon-inducible genes (*IFIT3, IFITM1, IFI44L, IFIT, IFITM8P* and *IFITM3*) are positively correlated with YAP1 expression in *CellMiner-Global*. By contrast, the NE subtypes show negative correlation between *NEUROD1* and *ASCL1* expression for those same immune genes (Figure 6G).

Based on the study of Wang et al. (2019) reporting a novel antigen presentation machinery transcription signature score (APM) yielding a high prediction index for tumor response to immune checkpoint inhibitors (ICI) in conjunction with tumor mutation burden (TMB), we tested the APM score in the SCLC cell lines (Supplementary Figure S13). The APM score showed a high correlation with *PD-L1* expression, which is notable as *PD-L1* is not included in the 13 genes constituting the APM score. Also, the SCLC-Y subtype showed the highest APM score (Supplementary Figure S13), consistent with the potential activation of their antigen presentation and innate immune response pathways.

### Cell Surface Biomarkers for Targeted Therapy in Relation with the NAPY Classification

Antibody-targeted therapies including antibody-drug conjugates (ADC) represent a promising approach for specific homing, increased uptake and drug retention at tumor sites while reducing drug exposure to normal tissues and the associated dose-limiting side effects (Coats et al., 2019). Proof of concept in SCLC has been established for Rovalpituzumab tesirine (Rova-T), the ADC targeting DLL3 with a DNA-crosslinking warhead (Das, 2017).

A primary criterium for efficient drug delivery treatment is to choose an exclusively or overexpressed target for the cancer cells. Figure 6I and Supplemental Figure S14 shows the expression of two receptors of clinical ADCs in the SCLC cell lines: DLL3 [used for SCLCs as rovalpituzumab tesirine (Morgensztern et al., 2019; Rudin et al., 2017)] and the carcinoembryonic antigen CEMC5 [used in other clinical indications as Labetuzumab govitecan (Das, 2017)]. Figure 6I shows that *DLL3* expression is highly correlated with *ASCL1* expression (Pearson correlation = 0.62), suggesting that treatments targeting DLL3, such as rovalpituzimab tesirine, could be selective toward SCLC-A tumors (Rudin et al., 2019). *CEACAM5* is highly expressed in only a subset of SCLC-A cell lines, which may be potentially sensitive to labetuzumab govitecan (IMMU-130) and other ADCs using CEACAM5 as their targeted receptor. Both DLL3 and CEACAM5 have their highest expression in SCLC among all GDSC tissue types (Supplemental Figure S14). Expression of *TACSTD2* (*TROP2)*, which is used as target for sacituzumab govitecan (IMMU-132) in patients with triple-negative breast cancer (TNBC), exhibits a low expression level in all SCLC cell lines, suggesting that using *TACSTD2* as targeted receptor may not be efficient in SCLC (Supplemental Figure S15).

Among potential new targets for the development of ADCs, the previously described specific neuroendocrine markers *NCAM1, CD24, CADM1* and *ALCAM* are highly expressed in non-YAP1 SCLC (Figure 6J), suggesting the potential of developing ADCs targeting such surface receptors for NE-SCLC and SCLC-P patients. In contrast, the non-neuroendocrine surface markers *CD151* and *EPH2* are highly expressed in the *YAP1* cell lines (Figure 6K), suggesting their potential as target receptors for SCLC-Y cancers.

## Discussion

SCLC CellMiner (https://discover.nci.nih.gov/SclcCellMinerCDB/) provides a unique resource including the most extensive SCLC datasets not only in terms of number of cell lines but also by its extensive omics and drug sensitivity databases. It also includes high resolution methylome data, which were performed for the purpose of the current study. SCLC CellMiner enables casual and experienced user to perform cross-comparison for all the omic and drug features of the SCLC cell lines of the NCI-DTP (SCLC NCI/DTP), Sanger-MGH (SCLC GDSC), Broad-MIT (SCLC CCLE and SCLC CTRP) and UT Southwestern (SCLC UTSW). It demonstrates the high reproducibility of the data for given cell lines across databases, which led to building an integrated platform (“SCLC Global”) to search genomic and drug features across the whole 116 cell line database.

Human cancer-derived cell lines remain the most widely used models and the primary basis to study the biology of cancers. They also enable the testing of new drugs and determinant of response hypotheses to improve cancer treatment (Gillet et al., 2013; Marx, 2014). A recent example is the discovery of SLFN11 as a dominant determinant of response to widely used chemotherapeutic agents targeting replication including topoisomerase inhibitors, platinum derivatives, gemcitabine and hydroxyurea as well as PARP inhibitors (Barretina et al., 2012; Murai et al., 2019; Zoppoli et al., 2012). Hence, the large database of SCLC cell lines offers a spectrum of models with the full genetic and molecular diversity seen in this subtype of cancer, as exemplified by the clear division of the 116 cell lines across the four recently proposed subgroups of SCLCs (NAPY classification) (Rudin et al., 2019). Although it appears that at the genomic level driver mutations are retained, several studies reveal a drift at the transcriptomic level, leading to the conclusion that cancer cell lines bear more resemblance to each other, regardless of the tissue of origin, than to the clinical samples that they are supposed to model. However, several other studies have come to the opposite conclusion, demonstrating the need for human cancer cell line panels (Barretina et al., 2012; Neve et al., 2006; Reinhold et al., 2019; Wang et al., 2006; Weinstein, 2012; Zoppoli et al., 2012). Although it was believed that tumor cells lost their differentiated properties during cell culture, it was later shown that this “dedifferentiation” was the result of stromal cell overgrowth and that “true” tumor cell cultures often retained their differentiated properties (Sato, 2008). For lung cancer cell lines, it has been shown that the genomic drift during culture life is not as great as commonly believed (Wistuba et al., 1999). The recent analyses across SCLC cell lines, PDX models and human tissues reported by Rudin et al. (2019) and our present analyses support this conclusion.

SCLC is known to be highly proliferative (Gazdar et al., 2017) and to be under replication stress (Thomas and Pommier, 2016). The SCLC CellMiner transcriptome data provide evidence confirming that specific feature. Indeed, genes known to be involved in DNA replication exemplified by *PCNA, MKI67* (encoding Ki67), *FEN1* and *PARP1* are highly expressed in SCLC compared to the other subtypes of cancers (Supplemental Figure S16). Moreover, we find evidence of chromatin alteration in SCLC. Not only are many core histone genes hypermethylated (see Figure 3) but also *H2AFY*, a non-canonical histone belonging to the H2A family encoding macroH2A.1, exhibits high expression in the SCLC cell lines. Two H2AFY splice variants have been identified and SCLC cell lines predominantly express high levels of the macroH2A1.2 variant compared to macroH2A1.1 (both encoded by *H2AFY*). The macroH2A1.2 splice variant is known to promote homologous recombination and is essential for proliferation (Kim et al., 2018). This further underscores the highly proliferative characteristic of SCLC cell lines, in addition to the overexpression of the MYCs genes (see Figure 4 and Supplementary Figure S2 and S6).

In the context of chromatin and the histone genes, *ACTL6B*, which encodes a subunit of the BAF (BRG1/brm-associated factor) complex in mammals is highly expressed in the SCLC cell lines (Supplemental Figure S17). The BAF complex is functionally related to SWI/SNF complexes, which are known to facilitate transcriptional activation of specific genes by antagonizing chromatin-mediated transcriptional repression. Interestingly, we found that the expression of *ACTL6B* is high and specific to SCLC and brain tumor cell lines and that its expression is highly correlated with other the expression of other chromatin genes including *HMGN2, KDM4B* and *SMARCA4* (Supplemental Figure S17). Among the SCLC cell lines, only the neuroendocrine cell lines (high *ASCL1* or high *NEUROD1*) harbor high expression of *ACTL6B* while the *YAP1* SCLC cell lines express significantly less *KDM4B* and *SMARCA4* (Supplemental Figure S17). These results suggest that this specific BAF complex subunit is critical in neuroendocrine SCLCs.

Supporting the importance of epigenetics in SCLC carcinogenesis, we provide an extensive DNA methylation database including the methylome of 66 cell lines from the NCI performed by high resolution Affymetrix 850k array and the analysis of 61 cell lines from the GDSC analyzed by 450k Array (see Figures 1 and 3) and demonstrate that SCLC cell lines exhibit a distinct methylation profile. First, they are globally hypomethylated, suggesting a plasticity of SCLC cell lines compared to the other cancers. Secondly, they exhibit a distinct and coherent profile of methylation compared with other subtypes of cancers, especially NSCLC (see Figure 3). Interestingly, most of genes with low methylation are involved in neurological pathway suggesting that neuroendocrine differentiation could be driven by epigenetic and especially DNA promoter methylation. Only a few studies focused on SCLC methylation profile. In 2013, Kalari *et al*. found consistent results and identified more than one hundred specifically hypermethylated genes in SCLC with gene ontology analysis indicating a significant enrichment of genes involved in neuronal differentiation (Kalari et al., 2013). By contrast, Poirier et al. (2015) reported that SCLC tend to have a high methylation level. The apparent discrepancy could be due to the fact that they included PDX and tumor samples and that they did not measure the global level of promoter methylation, as we have done, but the proportion of highly variable CpGs. Yet, they concluded, that high methylation instability is consistent with the plasticity of SCLC (Poirier et al., 2015).

SCLC CellMiner validates the recently proposed SCLC NAPY classification (Rudin et al., 2019) (see Figure 4), and provides insights into the four NAPY genes and their coordinated pathway network and connections with the NOTCH pathway (Figures 5). The coregulation of many functionally related genes is notable for the ASCL1 and YAP1 pathways examined in Figure 5. Indeed, *ASCL1* expression is highly correlated with the expression of its transcription coactivators *NKX2-1* and *PROX1* in spite of their different chromosome locations. The same observation applies to the YAP1/TAZ (WWTR1) heterodimer, suggesting master regulators upstream from the *ASCL1* and *YAP1* genes. Identifying those potential regulators (super-enhancers, microRNAs or non-coding RNAs) warrants further investigations, which hopefully will be fostered by the SCLC CellMiner resources. Unexpectedly, we found that the expression of the genes encoding the Hippo pathways (*MOB1A/B* and *LATS1/2*) and its coactivator (*AMOT* and *AMOTL2*) are co-expressed with highly significant correlation with *YAP1*. This finding suggest that the SCLC-Y cell lines are primed with a potential negative feedback from the Hippo pathway. Consistent with the results of Rudin et al. (2019) *al*., the NAPY classification shows that the cell lines driven by *ASCL1* and *NEUROD1* often overlap (see Figure 4F) except for their relationship with the NOTCH pathway where the SCLC-A cells show a stronger negative correlation with NOTCH gene expression than the SCLC-N cells (see Supplementary Figure 9). Both *ASCL1* and *NEUROD1* are transcriptional regulators and main drivers of neuroendocrine pathways and the cell lines co-expressing both gene share common features in terms of co-expressed neuroendocrine genes, MYCL-MYCN overexpression, drug sensitivities and cell surface markers (see Figures 4 & 6), questioning how these two groups define clearly distinct entities.

Transcriptome and drug response analyses highlight the distinguishing features of the SCLC-Y cell lines. Indeed, by contrast to the three other transcription factors (ASCL1, NEUROD1 and POU2F3), *YAP1* expression is not specific to SCLC and *YAP1* is widely and differentially expressed across a wide range of cancer cell lines (see Figure 4) (Ma et al., 2019). Notably, transcriptome analyses cluster the SCLC-Y with NSCLC cell lines, suggesting a different cellular origin for the SCLC-Y cancers (see Figure 5F). The SCLC-Y cell lines also express the NOTCH pathway, which is opposite to the SCLC-A neuroendocrine cell lines (see Figure 5 and Supplementary Figure S9). This differential feature could be related to the direct transcriptional activation of the NOTCH pathway by YAP/TAZ (see Figure 5C) (Yimlamai et al., 2014). In addition, SCLC-Y cell lines do not express *MYCL* or *MYCN* but rather *MYC* (see Figure 4), and consistent with the results of McColl et al. (2017), SCLC-Y cell lines tend and not to be mutated for *RB1* (see Figure 5H) and to express *RB1*, which is not the case for the 3 other SCLC subtypes (see Figure S11). We also found that the SCLC-Y cells express the DNA replication and proliferation genes to a lower level than the other SCLC subgroups (see Supplemental Figures S11 & S16). Finally, the SCLC-Y cell lines were often derived from non-smoker patients (Supplementary Table S1 & Figure 18). One of the limitations of this finding is that many cell lines were not annotated, so these results concerning tobacco status require confirmation in a larger cohort. In total, our data highlight that SCLC-Y cell lines are probably derived from a different cell type compared to the other neuroendocrine SCLC.

The SCLC-Y also differ from the other subgroups, SCLC-N, A & P in terms of drug sensitivity. As demonstrated in Figures 6 & S12, while the SCLC-P cell lines are consistently among the most sensitive NAPY subgroup to the standard of care treatments (etoposide, cisplatin and topotecan) and to the PARP inhibitor talazoparib, the SCLC-Y cells are most resistant to those treatments. The SCLC-N and -A show a wide range of responses to those classical chemotherapies with some cell lines highly responsive and some not. A significant determinant of response to those standard of care treatments is *SLFN11* expression (Murai et al., 2019), with a broad range of expression and approximately 40% of the 116 SCLC cell lines expressing no or very low *SLFN11* transcripts (see Figures 6F & S12). Another potential determinant of response is MGMT with approximately 33% of the 116 SCLC cell lines expressing no or very low *MGMT* transcripts (see S12D), which suggest the potential of using temozolomide in such tumors, especially in the case of brain metastases (Pietanza et al., 2018; Thomas et al., 2017).

In spite of the resistance of non-neuroendocrine (or variant) SCLC cells (SCLC-P and -Y subgroups) to the standard of care treatments (Gazdar et al., 1992), we find that those subgroups appear responsive to mTOR and AKT inhibitors (see Figure 6D-E). Our result is consistent with a recent study (Wooten et al., 2019) showing that non-neuroendocrine SCLC cell lines are sensitive to PI3K-AKT-mTOR, AURKA inhibitors and HSP90 inhibitors. Moreover, we found that the main difference between sensitive and non-sensitive cell lines is activation of the EIF2 pathway (see Figures 6 and S12), which is consistent with the PI3K-AKT-mTOR and MKI inhibitors sensitivity of SCLC-Y and SCLC-P. This hypothesis could open new therapeutic options in SCLC using translation-targeted drugs in development (Bastide and David, 2018; Sulima et al., 2017). Treatments targeting the mTOR pathway in SCLC patients have been evaluated or are in ongoing clinical trials. The results with monotherapy were not successful (Tarhini et al., 2010). Our findings suggest that better results might be obtained with appropriate patient selection.

Three final therapeutic insights can be derived from our study. First, the SCLC-Y cell lines are the only NAPY subgroup with antigen presenting and native immune predisposition (see Figure 6) while the neuroendocrine SCLC are among the most immune silent cancer cell lines based on their transcriptome profiles (see Figures 6G-H and S13). If verified in clinical samples, this finding might enable the selection of SCLC patient of the YAP1-expressing subgroup for immune checkpoint treatments. The second insight concerns the existence of potential surface markers that could be targeted selectively for the NAPY subgroups. As shown in the lower part of Figure 6, it is clear that the SCLC-Y cell lines express neither the therapeutically-relevant surface epitopes DLL3 or CEACAM5 (Das, 2017; Morgensztern et al., 2019; Rudin et al., 2017), which tend to be specific for the SCLC-A (and N) cancer cells. Yet, SCLC CellMiner could be used to identify potential surface markers of SCLC-Y cancers such as CD151 and EPHA2 (see Figure 6K). Finally, the SCLC-Y subgroup might respond to the YAP1 and NOTCH inhibitors in clinical development (Crawford et al., 2018; Leonetti et al., 2019).

Our analyses demonstrate the value of cancer cell line databases and imply that updating drug testing with new clinical drug candidates will provide valuable information to guide clinical trials. The results of our analyses also suggest the potential value of using the NAPY classification to select patients for targeted clinical trials. It is likely that genomic signatures based on genes expression (transcriptome) and DNA methylation (methylome) will have to be developed to build reliable tools to assign samples to each of the NAPY subgroups and determine their prognostic and therapeutic value. It also appears important to perform single-cell transcriptome and omic analyses, sequential biopsies and biopsies of different tumor sites to evaluate the tumor heterogeneity and plasticity of SCLCs.

## Material and methods

SCLC CellminerCDB is dedicated CellminerCDB version for SCLC cell lines (Reinhold et al., 2012; Reinhold et al., 2014; Reinhold et al., 2019; Reinhold et al., 2017b) https://discover.nci.nih.gov/cellminercdb/).

### SCLC-CellMinerCDB resources

The cell line sets included in SCLC-CellMiner Cross-Data-Base (CDB) currently are from the National Cancer Institute SCLC cell lines from the Developmental Therapeutics Program Small Cell Lung Cancer Project (SCLC NCI-DTP), Cancer Cell Line Encyclopedia (CCLE), Genomics and Drug Sensitivity in Cancer (GDSC), Cancer Therapeutics Response Portal (CTRP), the University of Texas SouthWestern (UTSW) and a new merge resource Global expression SCLC (add help section SCLC CellMiner CDB URL address). The data source details are described in “Help” section of the SCLC CellMiner website.

### SCLC-CellMinerCDB data

Most of the data including drug activity and genomics experiments were processed at the institute of origin and were downloaded from their website or provided from their principal investigator. However, methylation, mutation and copy number data were processed at Development Therapeutics Branch, CCR, NCI to generate a gene level summary as described previously (Barretina et al., 2012; Garnett et al., 2012; McMillan et al., 2018; Polley et al., 2016).

### DNA methylation data

Gene-level methylation using the 850k Illumina Infinium MethylationEPIC BeadChip array was summarized based on (Reinhold et al., 2017b). In short, methylation data were normalized using the minfi package using default parameters, where probe-level beta-values and detection p-values were calculated for each probe. This provided 866,091 methylation probe measurements. Methylation probe beta-values for individual cell lines with detection p-values >=10-3 were set to missing. Also probes with median p-value >= 10-6 were set to missing for all cells and removed from the analysis. Probe locations on the human genome (hg19 version) defined by Illumina was used for the analysis, annotating proximal gene transcripts and CpG islands. Probes were designated as category “1” or “2”, with category “1” considered to be most informative. Category “1” probes overlapped CpG islands and they overlapped either the TSS region within a 1.5kb distance, the first exon or 5’-UTR region. Additionally, probes on the upstream shore of a CpG island with a maximal distance of 200bp from the TSS were also included as category “1” probes. Category “2” probes were positioned either in the upstream- or downstream shore of a CpG island and overlapping the first exon, or on the downstream shore of CpG islands overlapping a 200bp region from the TSS, or in 5’-UTR. In case of genes with multiple transcript start sites, the transcript methylation with the most negative correlation to the gene level expression was used. The analysis resulted in gene-level methylation values for 23,202 genes.

### Copy number

Genome wide copy number for the cell lines was estimated from the methylation array data using the Chip Analysis Methylation Pipeline (*ChAMP*) (Tian et al., 2017) package. *ChAMP* returns lists of genomic segments with putative copy number estimates. However, the estimate is not valid for regions with high methylation detection p-values. For this reason, regions spanning more than 1kb with at least 5 probes with high detection p-values (p>0.05) were filtered out. The copy number estimates were set to missing for those areas. Gene level copy number (for n=25,568 genes) was calculated for each gene individually, by calculating the average estimate between the transcription start sites and transcription end sites.

### RNAseq data

The RNA-seq gene expression data from UTSW SCLC were obtained from analyses based on (McMillan et al., 2018). The raw data have been previously submitted to dbGaP (accession phs001823.v1.p1). For CCLE, the RNA-seq data was downloaded from the broad institute portal at https://portals.broadinstitute.org/ccle/data (version 2016-06-17)

### Global expression data

We also generate a new Global SCLC dataset using all combined cell lines, averaging gene expression based on z-scored gene expression from all resources: NCI SCLC, CCLE, CTRP, GDSC and UTSW. For each experiment, genes were scaled across all cell lines to create a z-score normalized dataset. The data sources have a mixture of microarray and RNA-seq gene expression. To test for removal of batch effects by gene scaling (z-score normalization), principal component analysis (Partek Genomics suite v7.17.1222) was performed on the raw (Fig.S3A) and normalized data (Fig.S3B) for CCLE microarray and RNA-seq datasets.

### Pathway level correlation of expression and DNA methylation

The correlation between methylation and gene expression for multiple functional categories was calculated based on genes in Supplementary Table S4. For each category, the median correlation of the related genes was calculated to identify potential categories of interest.

### Predictive power of DNA copy number and methylation on transcript expression

Testing the predictive power of DNA copy number and methylation on transcript expression was performed with linear regression analysis (as seen in Fig3G). For each of the 15,798 genes with all three forms of data available (transcript, methylation, and copy number levels) a linear regression model was fit, with both copy number and methylation as independent variables and transcript expression as the dependent variable. The model provided coefficients for the copy number and methylation that gave the lowest squared error between fitted values and true expression. We separated individual contributions of these two factors for gene expression prediction using the method of relative importance (Gromping, 2006), using the lmg method (Bacher, 1983) from the R package *relaimpo* to compute individual R2 values. Total (or combined) R2 is the summation of these two. Square roots of the R2 values were multiplied by the sign of the coefficients of the factors in the combined model to get the value of R.

### Cluster analysis

The methylation heatmap was created with the *ComplexHeatmap* (Gu et al., 2016) R package (version 1.20.0) using the kmeans clustering available in the *Heatmap()* function of the package. The number of reported clusters was selected based on cluster stability and biological significance.

SCLC cell lines groupings according NEUROD1, ASCL1, POU2F3 and YAP1 expression, MYC genes expression and neuroendocrine status defined by the Gazdar classification (Zhang et al., 2018) were done using the CIMminer tool from CellMiner (https://discover.nci.nih.gov/cimminer/oneMatrix.do). The used parameters were Euclidean distance method and complete linkage as cluster algorithm.

SCLC and NSCLC cell line grouping was performed with the gene expression data from the GDSC microarray dataset using the t-SNE algorithm in R (v3.5.1). The random seed was set to 1, the Euclidean distance of genes was calculated with the *dist()* function with default settings. The t-SNE grouping was calculated using the *Rtsne()* function from the Rtsne (van der Maaten, 2014) package (v0.15) using the calculated distance matrix, with perplexity set to 10, and 5k maximum iterations.

The NCI SCLC drug activity heatmap was generated using Partek Software. First, drugs with coefficient of variation less or equal to 0.09 were filtered out. Then the remaining data for the selected 134 drugs (from originally 527) across the 66 SCLC lines were clustered using the hierarchical method based on Euclidean distance and complete linkage.

### Gene set enrichment analysis

A preranked gene set enrichment analysis was run in R using the *clusterProfiler* (Yu et al., 2012) and *ReactomePA* (Yu and He, 2016) packages. Pathways with an adjusted p-value below 0.05 were considered as significantly enriched. Single sample gene set enrichment score (APM score) was computed using the R package GSVA (version 1.28.0).

### Statistical methods

Correlations, heatmaps, and histograms were generated mostly using The R Project for Statistical Computing. Some plots and analysis (such as the Kruskal Willis test) were generated using Partek Genomics suite v7.17.1222 (https://www.partek.com/partek-genomics-suite/) or using SCLC CellMinerCDB and CellMinerCDB (http://discover.nci.nih.gov/cellminercdb).

Wilcoxon rank-sum tests were used to test the difference between continuous variables such as drug sensitivity and gene expression according NAPY classification. We considered changes significant if p-values were below 0.05. In the figures, p-values below 0.00005 were summarized with four asterisks, p-values below 0.0005 were summarized with three asterisks, p-values below 0.005 were summarized with two asterisks and p-values below 0.05 were summarized with one asterisk.

## Data availability

All newely generated datasets have been deposited to the Gene Expression Omnibus (GEO, https://www.ncbi.nlm.nih.gov/geo/) under the accession number **GSE145156**.

## Data for reviewers

Data can be accessed at https://www.ncbi.nlm.nih.gov/geo/query/acc.cgi?acc=GSE145156 using the reviewer token “wnyxcukabfgnhet”.

