## Supplemental figures for "SCLC_CellMiner: Integrated Genomics and Therapeutics Predictors of Small Cell Lung Cancer Cell Lines based on their genomic signatures"

#### Supplementary Figures S1-S17

#### SCLC_CellMiner: Integrated Genomics and Therapeutics Predictors of Small Cell Lung Cancer Cell Lines based on their genomic signatures

Camille Tlemsani, Lorinc Pongor, Luc Girard, Nitin Roper, Fathi Elloumi, Sudhir Varma,
Augustin Luna, Vinodh N. Rajapakse, Sebastian Robin, Kurt W. Kohn, Julia Krushkal, Beverly A. Teicher,
Paul S. Meltzer, William C. Reinhold, John D. Minna, Anish Thomas and Yves Pommier

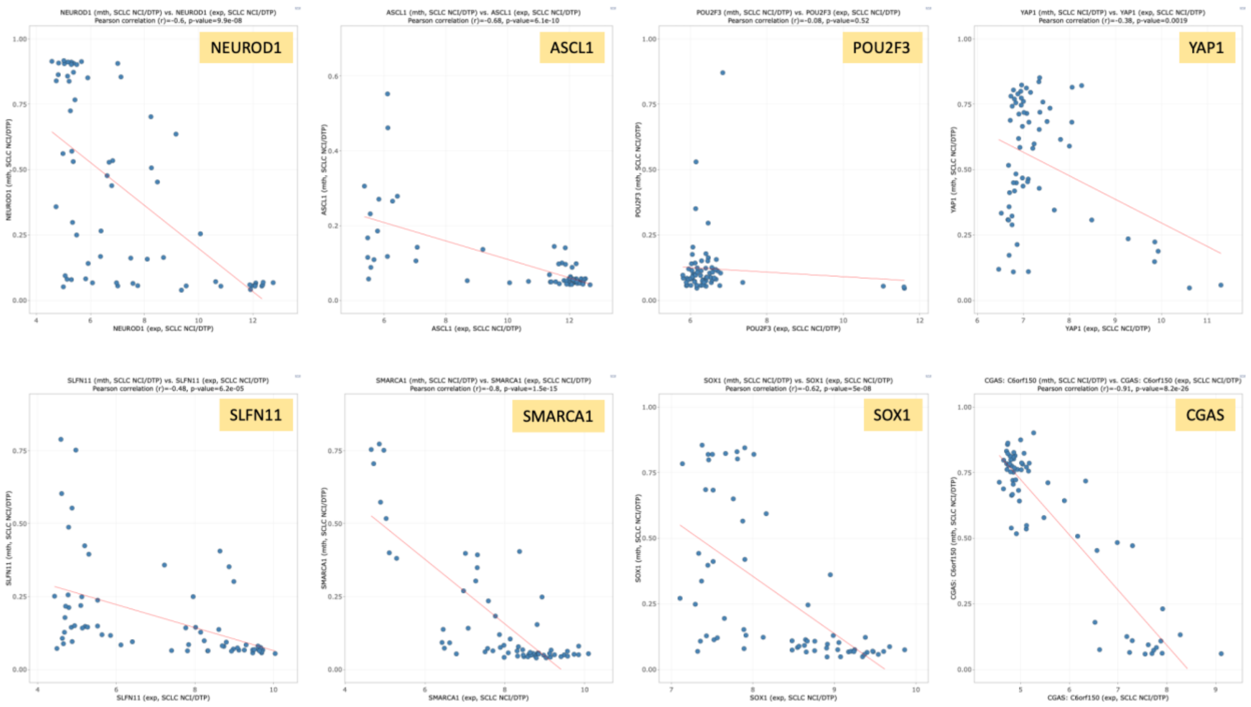


**Figure S1: Correlation between gene expression and promoter methylations for the NAPY genes (*NEUROD1, ASCL1, POU2F3 and YAP1*), *SLFN11*, *SMARCA1*, *SOX1* and *CGAS* in the SCLC cell lines**

Snapshot from SCLC_CellMiner (https://discover.nci.nih.gov/SclcCellMinerCDB) plotting DNA methylation (y-axis) vs gene expression (x-axis). The Pearson correlations of *NEUROD1*, *ASCL1*, *POU2F3*, *YAP1*, *SLFN11*, *SMARC1*, *SOX1* and *CGAS* genes in the NCI-SCLC dataset are -0.60, -0.68, -0.08, -0.38, -0.48, -0.80, -0.62 and -0.91, respectively.

Camille Tlemsani, Lorinc Pongor, Luc Girard, Nitin Roper, Fathi Elloumi, Sudhir Varma,
Augustin Luna, Vinodh N. Rajapakse, Sebastian Robin, Kurt W. Kohn, Julia Krushkal, Beverly A. Teicher,
Paul S. Meltzer, William C. Reinhold, John D. Minna, Anish Thomas and Yves Pommier

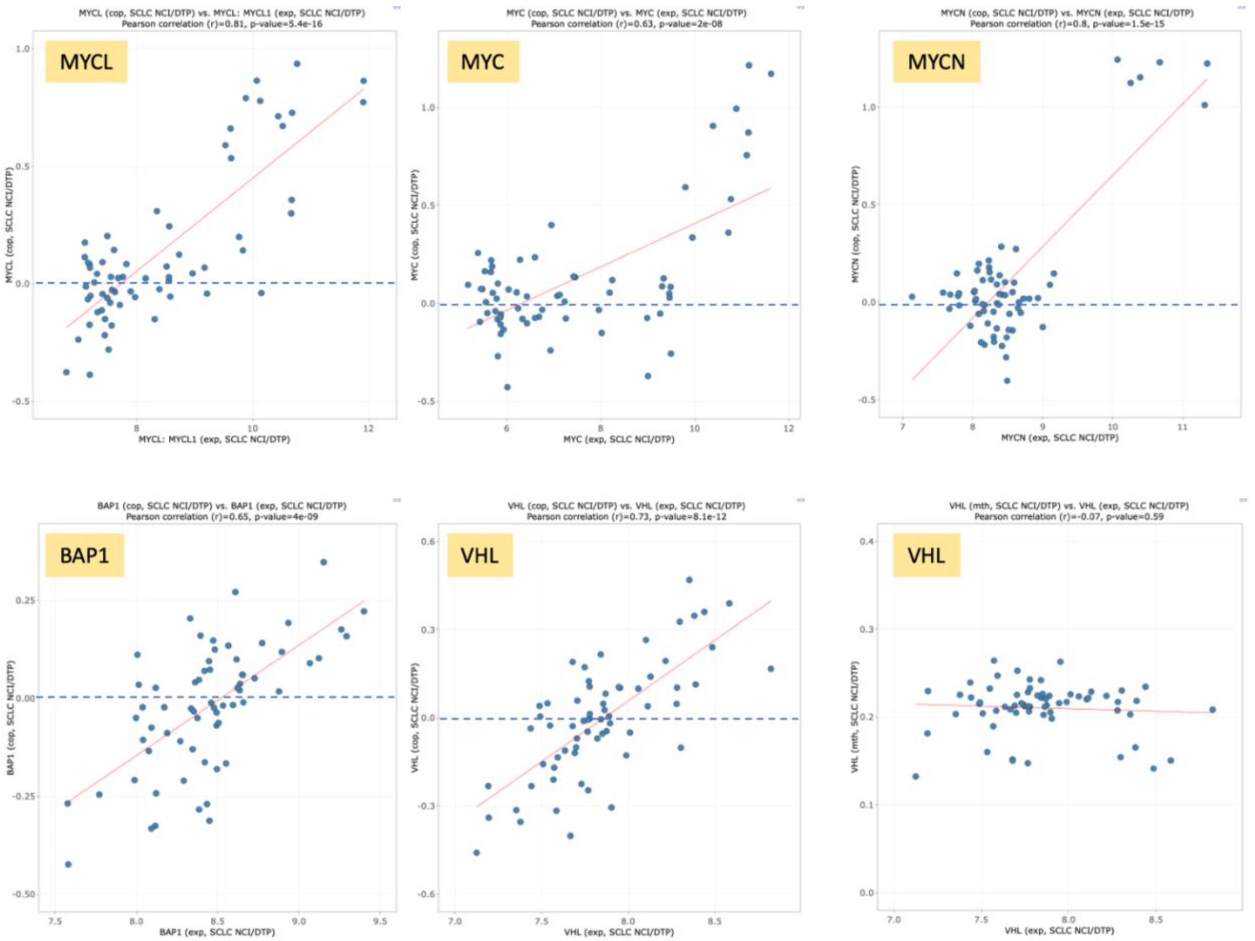


**Figure S2: Correlations between gene expression and gene copy number for *MYCL, MYC, MYCN, BAP1* and *VHL* genes**

Snapshot from SCLC_CellMiner (https://discover.nci.nih.gov/SclcCellMinerCDB) plotting gene copy number (y-axis) vs expression (x-axis). The Pearson correlations are 0.81, 0.63, 0.80, 0.65 and 0.73, respectively. The bottom right plot represents the methylation (y-axis) and expression (x-axis) for *VHL*. Pearson correlation - 0.07.

### SCLC_CellMiner: Integrated Genomics and Therapeutics Predictors of Small Cell Lung Cancer Cell Lines based on their genomic signatures

Camille Tlemsani, Lorinc Pongor, Luc Girard, Nitin Roper, Fathi Elloumi, Sudhir Varma, Augustin Luna, Vinodh N. Rajapakse, Sebastian Robin, Kurt W. Kohn, Julia Krushkal, Beverly A. Teicher, Paul S. Meltzer, William C. Reinhold, John D. Minna, Anish Thomas and Yves Pommier

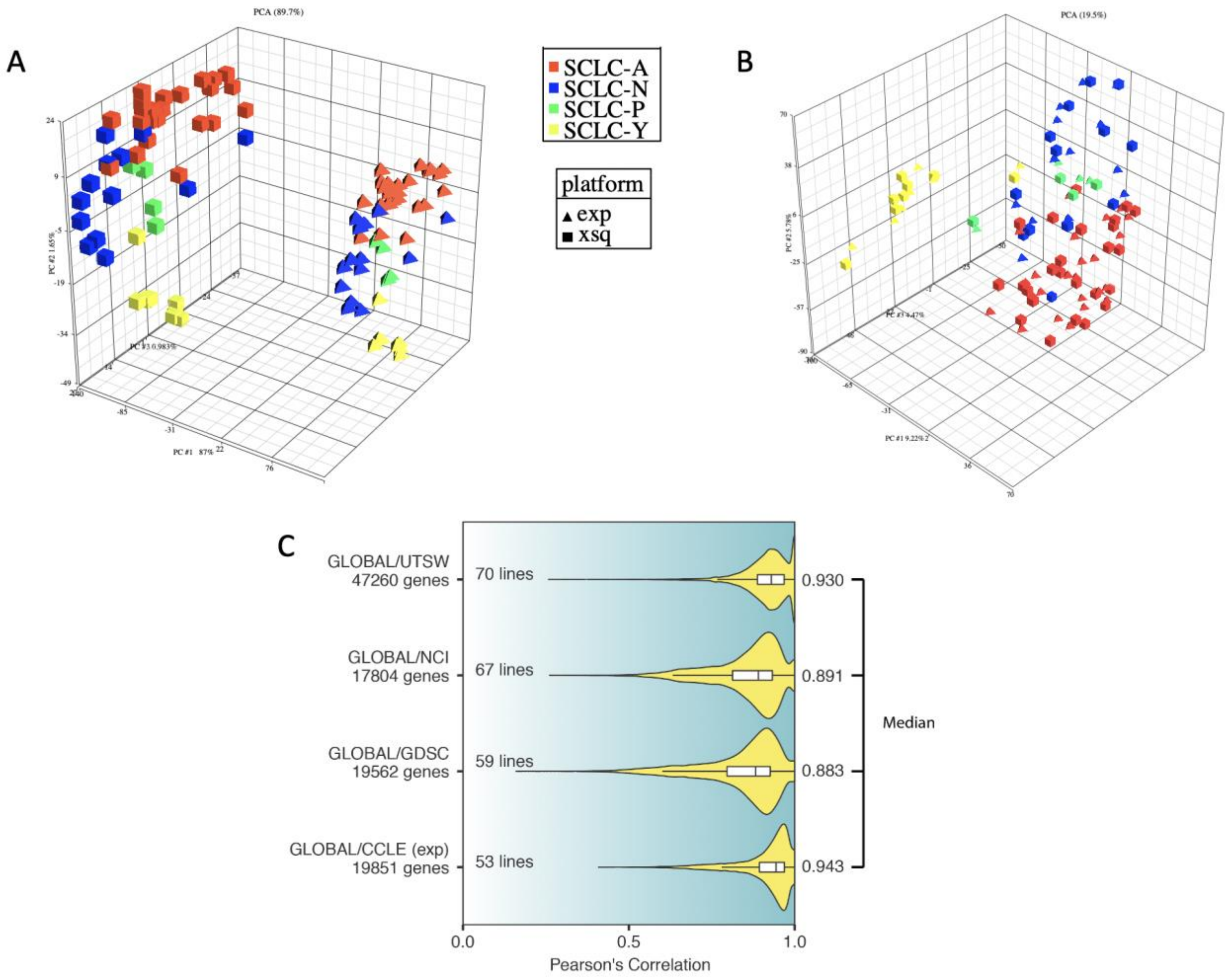


**Figure S3: Normalized expression data validation and reproducibility of the "Global" dataset (116 SCLC cell lines) using Z-score**

Principle component analysis (PCA) plots before (**A**) and after (**B**) Z-score normalization between microarray and RNAseq data in the CCLE dataset. Each point represents one cell line. Rectangles are the cell lines according RNAseq value and triangles the cell lines according microarray expression values. Cell lines are color-coded according to the NAPY classification: SCLC-A cell lines in red, SCLC-N cell lines in blue, SCLC-P in green and SCLC-Y in yellow The x-axis represents the first principle component, the y-axis the second component and the z-axis the third principle component. (**C**). Reproducibility between the Global dataset and the other datasets. Pearson correlations between the Global dataset and the indicated data sources for matched cell lines were 0.930, 0.891, 0.883 and 0.943 for GLOBAL/UTSW, GLOBAL/NCI, GLOBAL/GDSC and GLOBAL/CCLE (microarray) expressions, respectively.

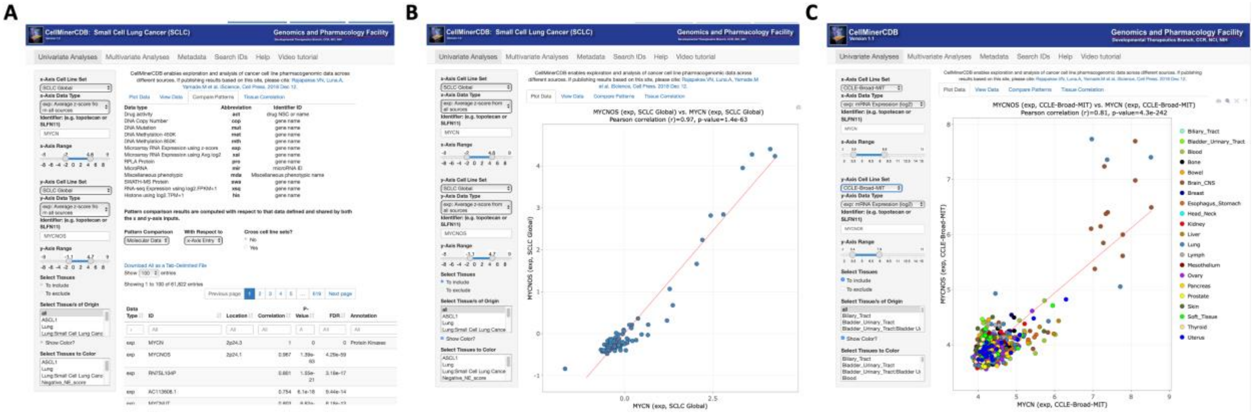


**Figure S4: SCLC-CellMinerCDB examples for "Compare Patterns" and correlation between *MYCN* and *MYCNOS* expression**

Each panel is a snapshot from SCLC-CellMiner and CellMiner website (https://discover.nci.nih.gov/SclcCellMinerCDB). (**A**) The correlation between *MYC* expression in all SCLC cell lines and all the other genes can be easily found through the "Compare Patterns" function in the "Univariate Analysis" section. This example shows that *MYC* expression is highly correlated with MYC antisense *MYCNOS* expression (Pearson correlation = 0.967, p-value =1.39x10$^{-63}$). (**B**) SCLC_CellMiner Snapshot showing *MYCNOS* expression (y-axis) vs *MYCN* expression (x-axis) in the SCLC_Global cell lines. (**C**) Findings in SCLC cell lines can be readily compared to other cancer cell lines using CellMinerCDB (http://discover.nci.nih.gov/cellminercdb). Shown is the high correlation between *MYCN* and *MYCNOS* expression is also present across additional cancer subtypes.

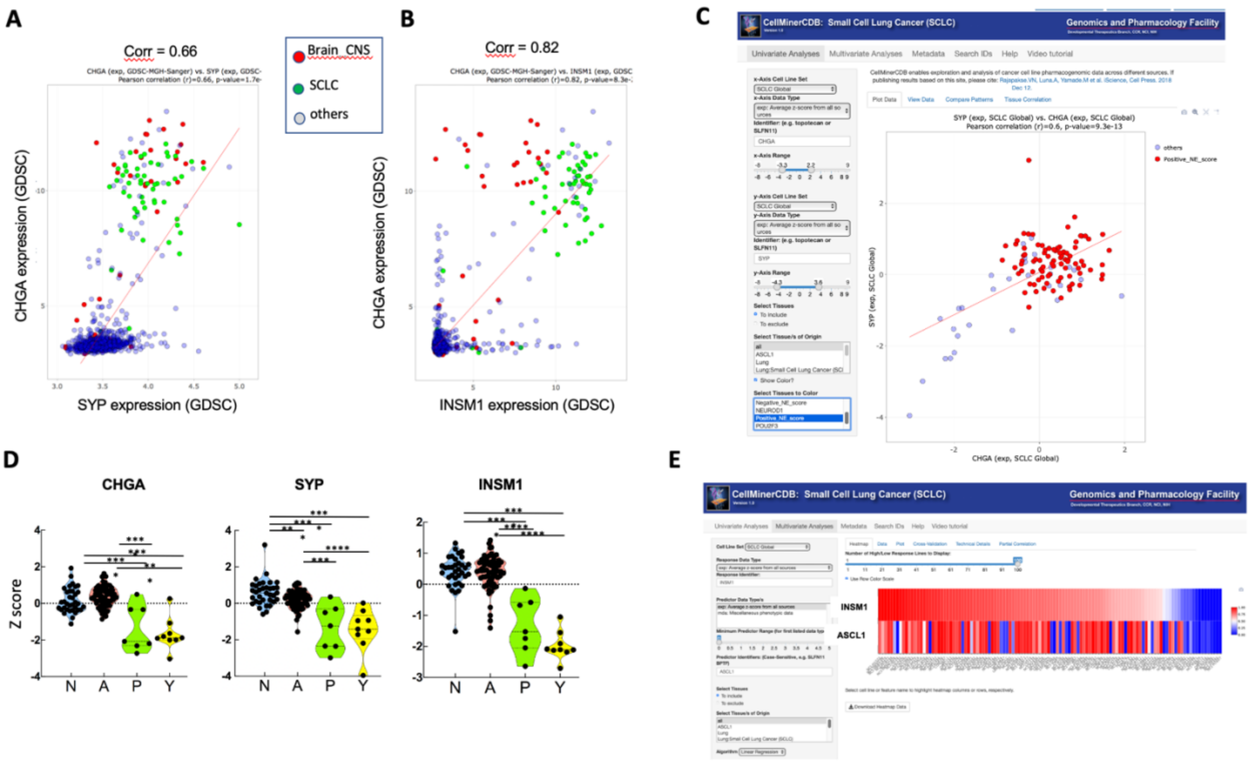


**Figure S5: CHGA, SYP and INSM1 neuroendocrine expression characteristics**

Chromogranin (CHGA), synaptophysin (SYP) and insulinoma-associated protein 1 (INSM1) are neuroendocrine markers used in routine practice to diagnose SCLC and NE tumors. The images for each panel are snapshots from the SCLC-CellMiner (https://discover.nci.nih.gov/SclcCellMinerCDB) and CellMiner (http://discover.nci.nih.gov/cellminercdb) websites. Panels (**A**) and (**B**) highlight that *CHGA*, *SYP* and *INSM1* expressions are highly correlated with Pearson correlations at 0.66 and 0.82, respectively. High expression of these three neuroendocrine markers is mainly found in SCLC (green points) and brain tumor (red points) cell lines. Using "Univariate Analysis" function in SCLC-CellMiner website, the plot in panel (**C**), showing SYP expression (y-axis) and CHGA expression (x-axis), highlights that high expression of both neuroendocrine markers is found only in the cell lines with a high NE score. Similarly, panel (**D**) shows the expression of *CHGA* (left), *SYP* (middle) and *INSM1* (right) according the NAPY classification. The expression of the three neuroendocrine markers is higher in the SCLC-N and -A cell lines (blue and red, respectively) compared to the SCLC-P and -Y cell lines (green and yellow, respectively). Each point represents a cell line. (**D**) Snapshot of the "Multivariate Analyses" tool of SCLC_CellMiner showing that *ASCL1* expression is highly associated with *INSM1*. Red and blue are cell lines with the highest and lower gene expression.

### SCLC_CellMiner: Integrated Genomics and Therapeutics Predictors of Small Cell Lung Cancer Cell Lines based on their genomic signatures

Camille Tlemsani, Lorinc Pongor, Luc Girard, Nitin Roper, Fathi Elloumi, Sudhir Varma,
Augustin Luna, Vinodh N. Rajapakse, Sebastian Robin, Kurt W. Kohn, Julia Krushkal, Beverly A. Teicher,
Paul S. Meltzer, William C. Reinhold, John D. Minna, Anish Thomas and Yves Pommier

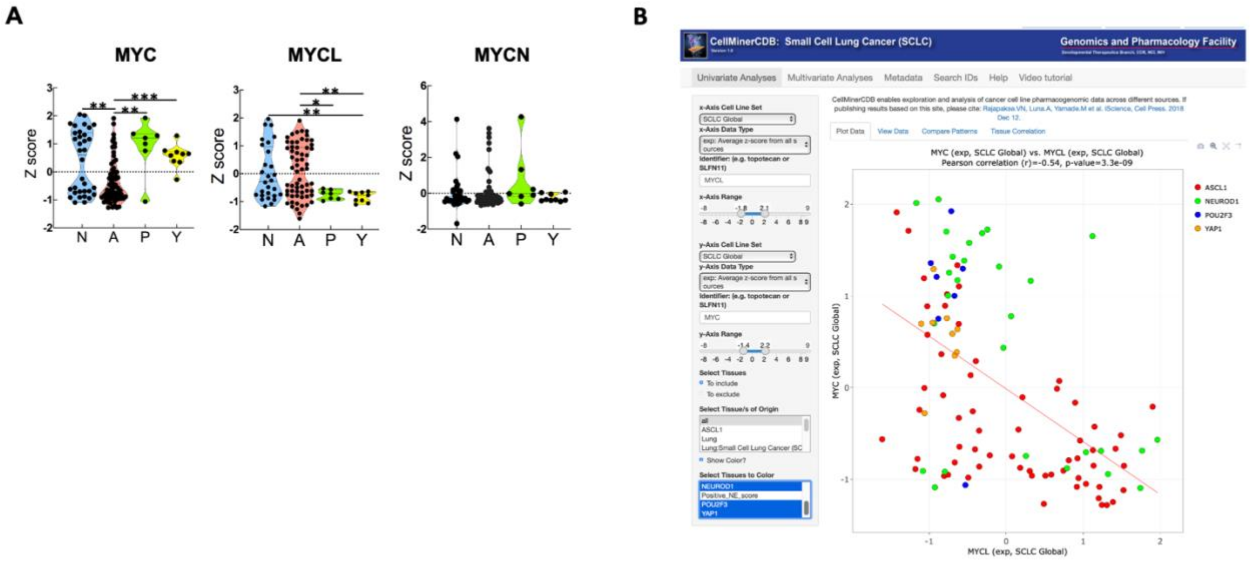


**Figure S6: Expression of the three MYC oncogenes according in relationship with the NAPY classification**

(**A**). Left: *MYC* is constantly highly expressed in the SCLC-P and -Y cell lines while its expression is bimodal in the SCLC-N cell lines and mainly under-expressed in the SCLC-Y cell lines. Middle: *MYCL* expression has an opposite behavior with low expression in all SCLC-P and -Y cell lines and a bimodal distribution in the SCLC-N and -A cell lines. Right: *MYCN* is less frequently expressed in all cell lines, especially the SCLC-Y subset. (**B**). Snapshot of *MYC* (y-axis) and *MYCL* expression (x-axis) in the 116 SCLC cell lines highlighting a high negative Pearson correlation between both genes (correlation = -0.54). High *MYC* expression is mainly found in the SCLC-P and -Y cell lines (blue and yellow points, respectively) while high *MYCL* expression is mainly found in the SCLC-A and -N cell lines (red and green points, respectively). The plot highlights that high *MYC* and *MYCL* expressions are mutually exclusive.

### SCLC_CellMiner: Integrated Genomics and Therapeutics Predictors of Small Cell Lung Cancer Cell Lines based on their genomic signatures

Camille Tlemsani, Lorinc Pongor, Luc Girard, Nitin Roper, Fathi Elloumi, Sudhir Varma,
Augustin Luna, Vinodh N. Rajapakse, Sebastian Robin, Kurt W. Kohn, Julia Krushkal, Beverly A. Teicher,
Paul S. Meltzer, William C. Reinhold, John D. Minna, Anish Thomas and Yves Pommier

A

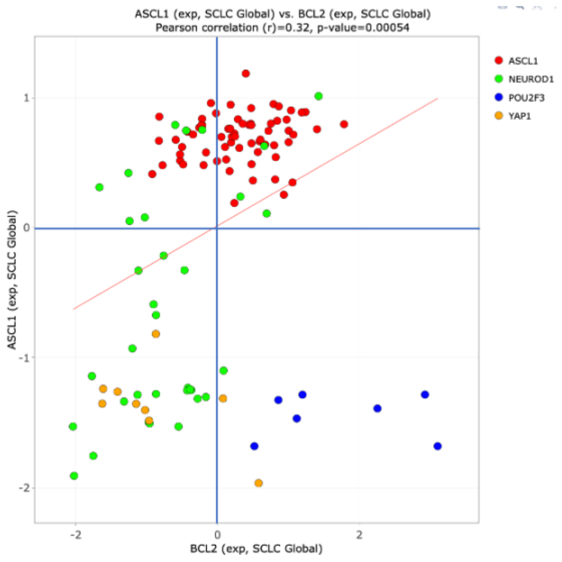


C

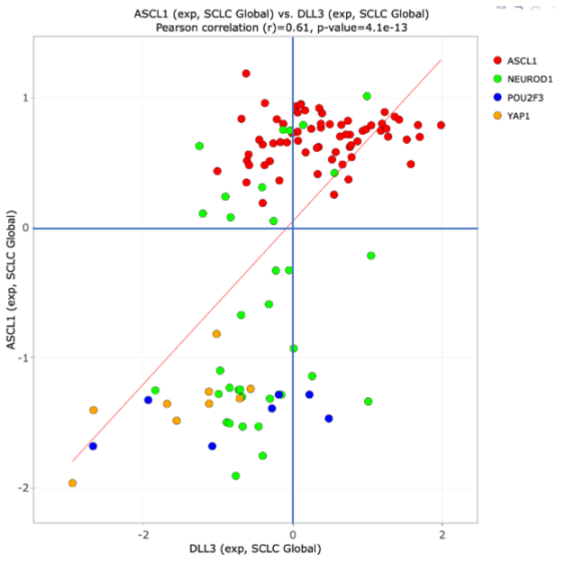


B

| | BCL2 (r) | P-value |
|---|---|---|
| N | -0.452 | 3.54e-7 |
| A | 0.316 | 0.000541 |
| P | 0.43 | 4.14e-6 |
| Y | -0.165 | 0.0769 |

D

| | DLL3 (r) | P-value |
|---|---|---|
| N | -0.176 | 0.0585 |
| A | 0.609 | 4.05e-13 |
| P | 0.277 | 0.00258 |
| Y | -0.54 | 4.07e-10 |

**Figure S7: *BCL2* and *DLL3* are highly expressed in the SCLC-A cell lines**

Correlations between *ASCL1* expression (y-axis) with *BCL2* (**A**) and *DLL3* (**C**) (x-axis) in the 116 cell lines of SCLC_Global. High *BCL2* and *DLL3* expressions are mainly found in the SCLC-A cell lines (red points). Of note, SCLC-P cell lines also have high *BLC2* expression. Details of Pearson correlations of each NAPY gene (*NEUROD1*, *ASCL1*, *POU2F3* and *YAP1*) vs *BLCL2* (**B**) and *DLL3* (**D**). The values in red represent the significantly positive Pearson correlations and the values in blue the significantly negative correlations.

### SCLC_CellMiner: Integrated Genomics and Therapeutics Predictors of Small Cell Lung Cancer Cell Lines based on their genomic signatures

Camille Tlemsani, Lorinc Pongor, Luc Girard, Nitin Roper, Fathi Elloumi, Sudhir Varma, Augustin Luna, Vinodh N. Rajapakse, Sebastian Robin, Kurt W. Kohn, Julia Krushkal, Beverly A. Teicher, Paul S. Meltzer, William C. Reinhold, John D. Minna, Anish Thomas and Yves Pommier

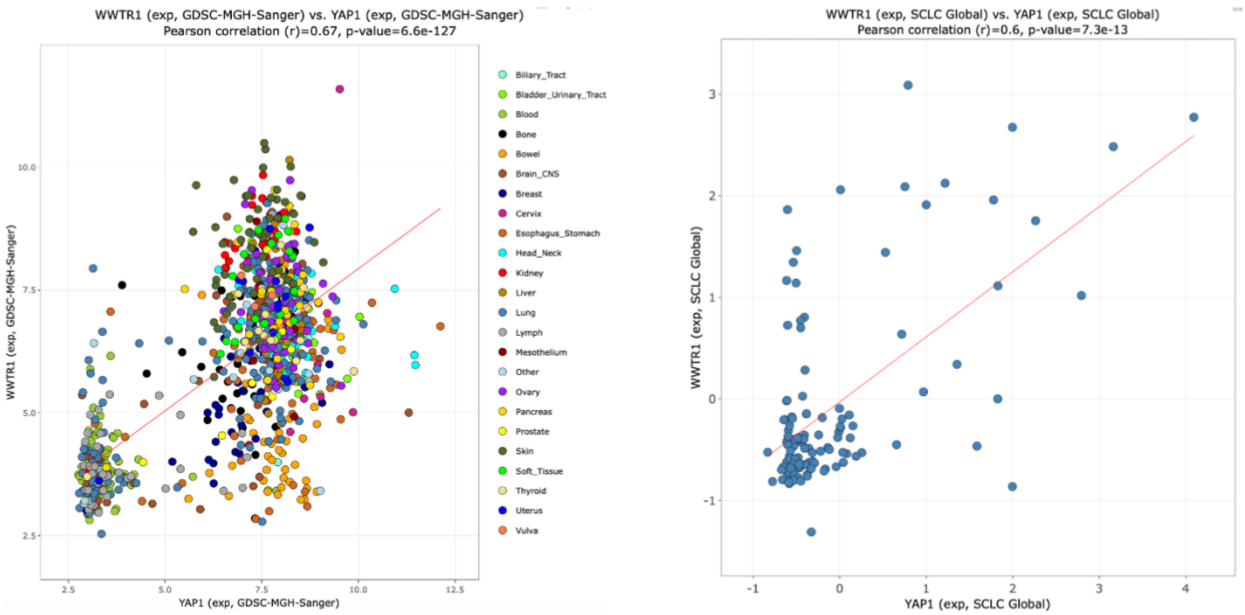


#### Figure S8: YAP1 and WWTR1 co-expression

Snapshot (http://discover.nci.nih.gov/cellminercdb; https://discover.nci.nih.gov/SclcCellMinerCDB) showing that *YAP1* expression is highly correlated with the expression of its heterodimeric partner TAZ (encoded by the *WWTR1/TAZ* gene) across the 986 cell lines of the GDSC (left) and among the 116 SCLC cell lines of CellMiner_Global (right).

### SCLC_CellMiner: Integrated Genomics and Therapeutics Predictors of Small Cell Lung Cancer Cell Lines based on their genomic signatures

Camille Tlemsani, Lorinc Pongor, Luc Girard, Nitin Roper, Fathi Elloumi, Sudhir Varma, Augustin Luna, Vinodh N. Rajapakse, Sebastian Robin, Kurt W. Kohn, Julia Krushkal, Beverly A. Teicher, Paul S. Meltzer, William C. Reinhold, John D. Minna, Anish Thomas and Yves Pommier

**A**

| | N | A | P | Y |
|---|---|---|---|---|
| NOTCH1 | -0.002 | -0.49 (1.84e-8) | 0.45 (4.62e-7) | 0.39 (1.68e-5) |
| NOTCH2 | -0.132 | -0.42 (3.07e-6) | 0.23 (1.49e-2) | 0.56 (6.96e-11) |
| NOTCH3 | -0.173 | -0.40 (8.55e-6) | 0.44 (8.96e-7) | 0.62 (2e-13) |
| NOTCH4 | -0.002 | 0.25 (0.00784) | -0.169 | -0.091 |
| REST | 0.221 (0.0171) | -0.55 (5.48e-10) | 0.51 (8.08e-7) | 0.76 (2.05e-19) |

**B**

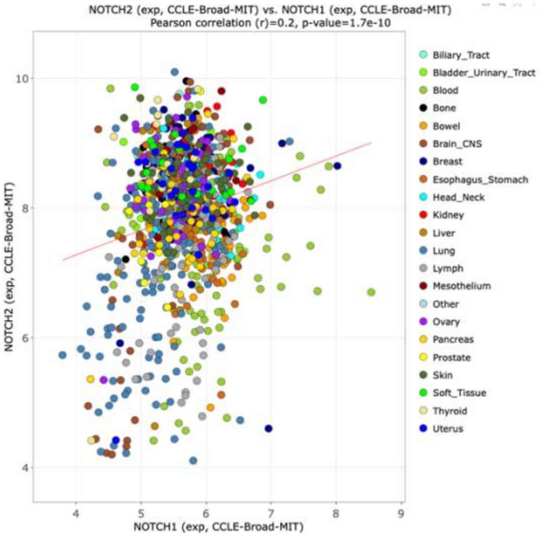


**C**

| Tissue of origin | Cell lines with complete observations | Pearson correlation | P-value |
|---|---|---|---|
| ALL | 1036 | 0.2 | 1.7e-10 |
| Brain_CNS | 86 | 0.58 | 5e-9 |
| Lung | 186 | 0.39 | 2.6e-8 |
| Lymph | 64 | 0.36 | 0.0038 |
| Thyroid | 12 | 0.64 | 0.026 |
| Pancreas | 44 | 0.32 | 0.037 |
| Esophagus_Stomach | 64 | -0.25 | 0.044 |
| Uterus | 27 | 0.32 | 0.11 |
| Prostate | 8 | 0.58 | 0.14 |
| Mesothelium | 11 | 0.45 | 0.16 |
| Ovary | 52 | 0.19 | 0.17 |
| Biliary_Tract | 8 | -0.45 | 0.27 |
| Bowel | 62 | -0.11 | 0.39 |
| Bladder_Urinary_Tract | 27 | -0.15 | 0.46 |
| Soft_Tissue | 20 | -0.12 | 0.61 |
| Blood | 116 | -0.044 | 0.64 |
| Bone | 29 | -0.086 | 0.66 |
| Breast | 59 | -0.035 | 0.79 |
| Head_Neck | 34 | -0.045 | 0.8 |
| Kidney | 36 | 0.04 | 0.82 |
| Liver | 28 | 0.03 | 0.88 |
| Skin | 62 | 0.016 | 0.9 |

#### Figure S9: NOTCH pathway transcriptional network in relationship with the NAPY classification

(**A**) Pearson correlations between the NOTCH/REST pathway gene expression and the NAPY genes (*NEUROD1*, *ASCL1*, *POU2F3* and *YAP1*). Expression of the NOTCH genes (except *NOTCH4*) is negatively correlated with *ASCL1* expression and positively correlated with *POU2F3* and *YAP1* expressions. (**B**) Snapshot from CellMinerCDB (http://discover.nci.nih.gov/cellminercdb) showing correlated expression of *NOTCH2* (y-axis) and *NOTCH1* (x-axis) across histological cell line subtypes in the CCLE dataset. (**C**) Snapshot of the tabular output from CellMinerCDB showing Pearson correlations between *NOTCH2* and *NOTCH3* expressions across cell lines from different histological subtypes.

**SCLC_CellMiner: Integrated Genomics and Therapeutics Predictors of Small Cell Lung Cancer Cell Lines based on their genomic signatures**

Camille Tlemsani, Lorinc Pongor, Luc Girard, Nitin Roper, Fathi Elloumi, Sudhir Varma, Augustin Luna, Vinodh N. Rajapakse, Sebastian Robin, Kurt W. Kohn, Julia Krushkal, Beverly A. Teicher, Paul S. Meltzer, William C. Reinhold, John D. Minna, Anish Thomas and Yves Pommier

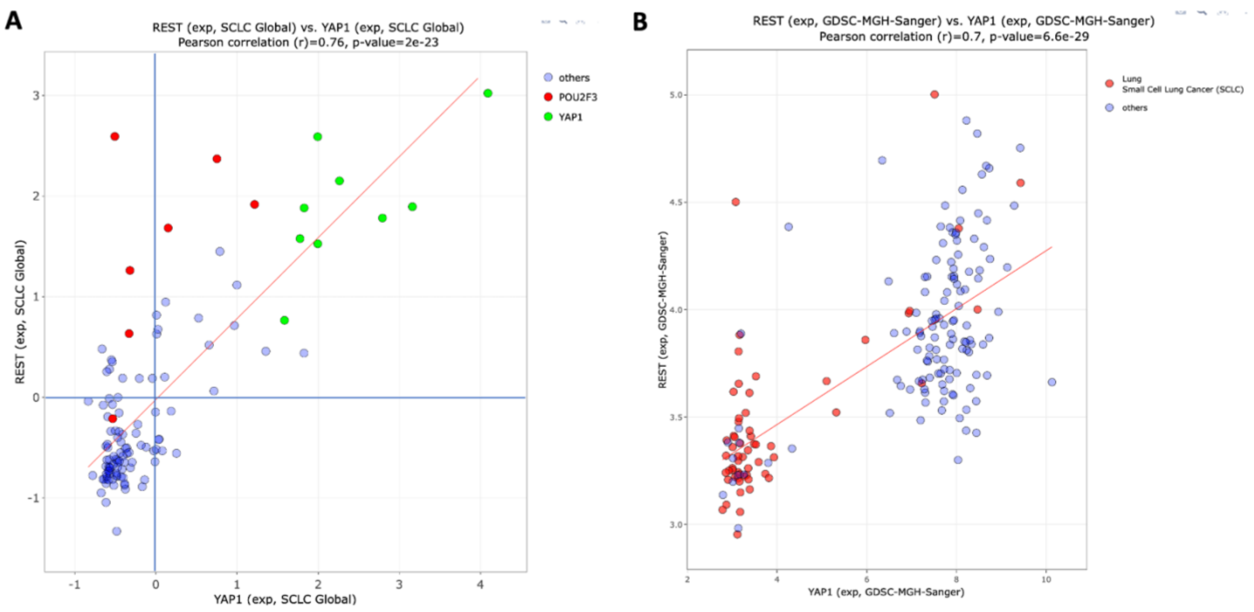


**Figure S10: REST expression as surrogate of NOTCH activation in the non-neuroendocrine SCLC cell lines SCLC-Y and SCLC-P**

Snapshots from SCLC_CellMiner showing *REST* (y-axis) vs *YAP1* (x-axis) expression in the 116 SCLC cell lines of SCLC_Global (http://discover.nci.nih.gov/cellminercdb) (**A**) and in all subtypes of lung cancers (SCLC + NSCLC) in the GDSC dataset (https://discover.nci.nih.gov/SclcCellMinerCDB) (**B**). The plots highlight that all non-neuroendocrine cell lines have a high *REST* expression (green and red point in panel (**A**)) and that high *YAP1* expressing lung cancer cell lines (**B**) also have high expression of *REST*.

**SCLC_CellMiner: Integrated Genomics and Therapeutics Predictors of Small Cell Lung Cancer Cell Lines based on their genomic signatures**

Camille Tlemsani, Lorinc Pongor, Luc Girard, Nitin Roper, Fathi Elloumi, Sudhir Varma,
Augustin Luna, Vinodh N. Rajapakse, Sebastian Robin, Kurt W. Kohn, Julia Krushkal, Beverly A. Teicher,
Paul S. Meltzer, William C. Reinhold, John D. Minna, Anish Thomas and Yves Pommier

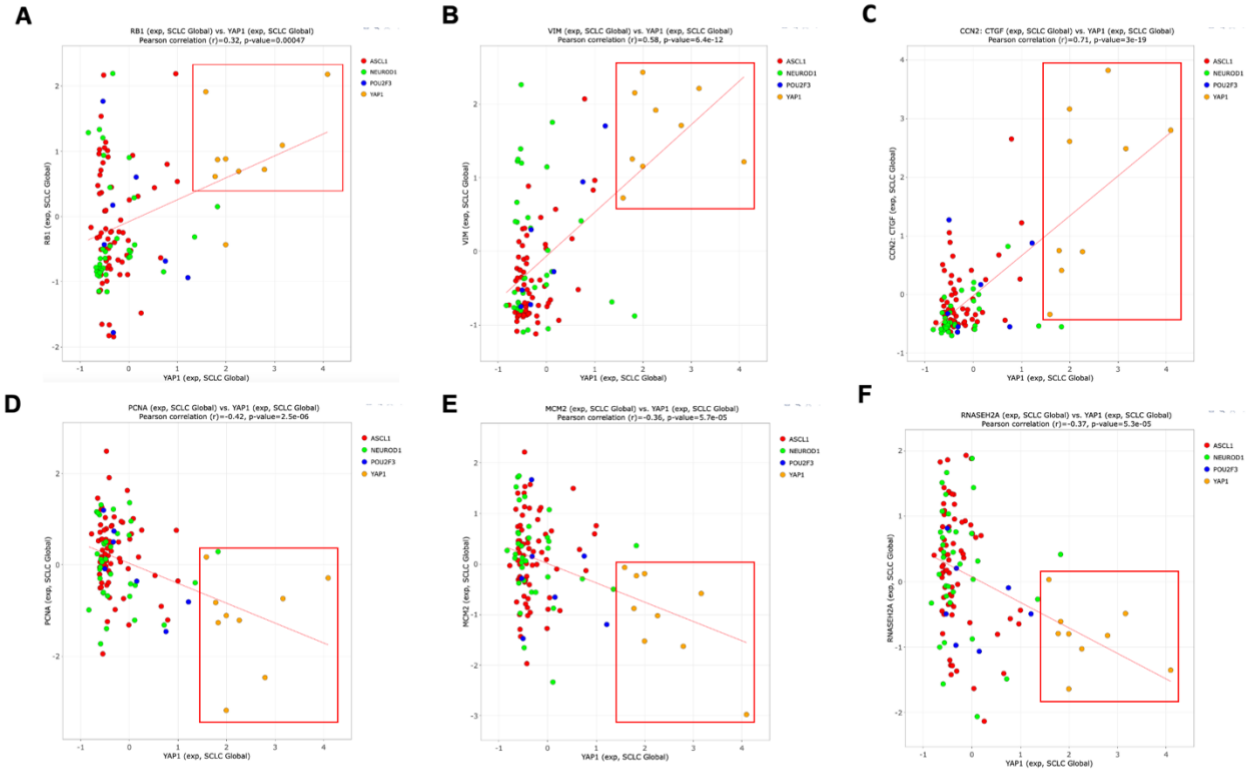


**Figure S11: Examples of correlations between the expression of *YAP1* and key cancer genes**

Snapshots (https://discover.nci.nih.gov/SclcCellMinerCDB) of *YAP1* expression (x-axis) vs the indicated cancer genes (y-axis) in the 116 cell lines of SCLC_Global. Each point represents a cell line (red: SCLC-A, green: SCLC-N, blue: SCLC-P and yellow: SCLC-Y). Panels (**A**), (**B**) and (**C**) show highly positive Pearson correlations between *YAP1* and *RB1*, *VIM* and *CCN2*. Panels (**D**), (**E**) and (**F**) show highly negative Pearson correlations between *YAP1* and the replication-associated genes *PCNA*, *MCM2* and *RNASEH2A*. The red rectangles encompass the SCLC-Y cell lines.

Camille Tlemsani, Lorinc Pongor, Luc Girard, Nitin Roper, Fathi Elloumi, Sudhir Varma, Augustin Luna, Vinodh N. Rajapakse, Sebastian Robin, Kurt W. Kohn, Julia Krushkal, Beverly A. Teicher, Paul S. Meltzer, William C. Reinhold, John D. Minna, Anish Thomas and Yves Pommier

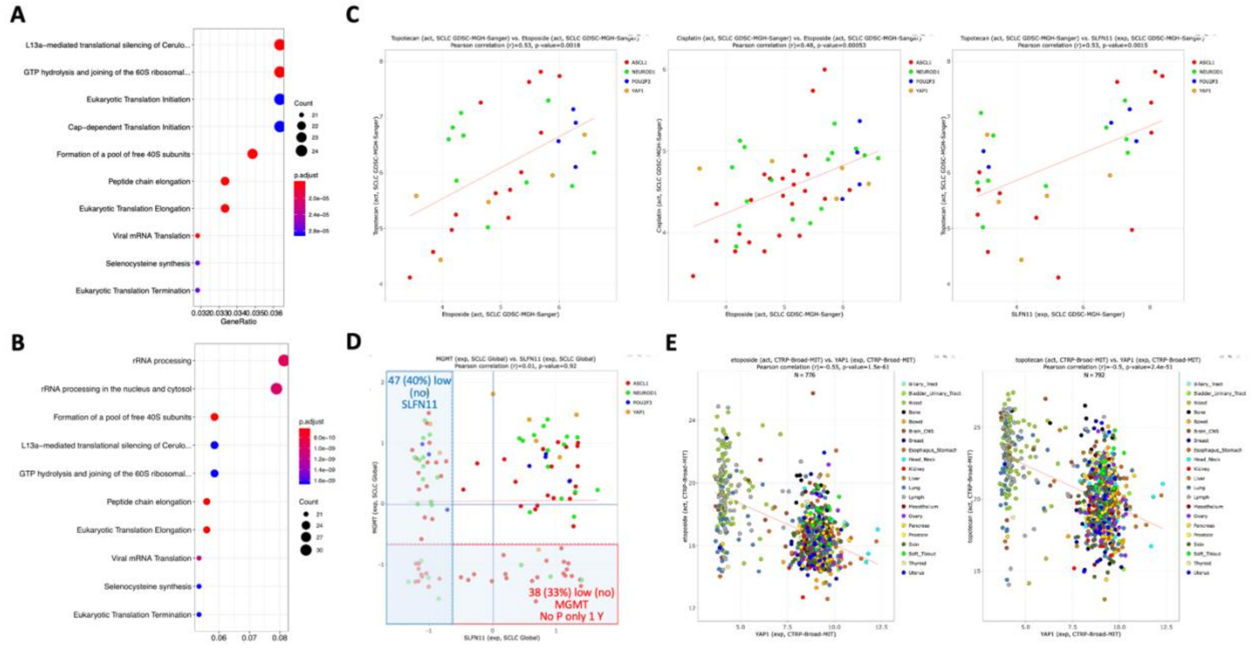


#### Figure S12: Drug sensitivity of the SCLC cell lines

(**A-B**) Pathway analyses for the differentially expressed genes between sensitive and non-sensitive cell lines according the heatmap analysis showed in Figure 6A. Panel (**A**) shows the pathway analysis when considering all genes and panel (**B**) the pathway analysis when considering the genes overexpressed in the most sensitive cell lines group according to Figure 6A. (**C**) Left: correlation between the activity of etoposide (-log IC50) (x-axis) and topotecan (y-axis) across the SCLC cell lines. Middle: correlation between the activity of etoposide (-log IC50) (x-axis) and cisplatin (y-axis). Right: correlation between *SLFN11* expression (x-axis) and the activity of topotecan (y-axis) in the SCLC cell lines. Each point represents a cell line (red: SCLC-A cell lines, green: SCLC-N cell lines, blue: SCLC-P cell lines and orange: SCLC-Y cell lines). (**D**) Broad range of *SLFN11* and *MGMT* expression in the 116 cell lines of SCLC_Global. Forty percent of the cell lines (47/116) have a low expression of *SLFN11* and 33% (38/116) low *MGMT* expression. Among them, there is no SCLC-P cell line and only one SCLC-Y cell line. (**E**) *YAP1* expression (x-axis) is negatively correlated with the activity of etoposide (log -IC50) (left plot) and also negatively correlated with the activity of topotecan (right) across different histological subtypes (http://discover.nci.nih.gov/cellminercdb).

### SCLC_CellMiner: Integrated Genomics and Therapeutics Predictors of Small Cell Lung Cancer Cell Lines based on their genomic signatures

Camille Tlemsani, Lorinc Pongor, Luc Girard, Nitin Roper, Fathi Elloumi, Sudhir Varma, Augustin Luna, Vinodh N. Rajapakse, Sebastian Robin, Kurt W. Kohn, Julia Krushkal, Beverly A. Teicher, Paul S. Meltzer, William C. Reinhold, John D. Minna, Anish Thomas and Yves Pommier

**A**

18 gene signature
https://elifesciences.org/articles/49020

| Protein Name | Gene Name | chromosome |
|---|---|---|
| B2M | B2M | 15q21-q22.2 |
| calreticulin | CALR | 19p13.3-p13.2 |
| Calnexin | CANX | 5q35 |
| ERAP1 | ERAP1 | 5q15 |
| ERAP2 | ERAP2 | 5q15 |
| HLA I | HLA-A | |
| HLA I | HLA-B | 6p21.3 |
| HLA I | HLA-C | 6p21.3 |
| ERp57 | PDIA3 | 15q15 |
| LMP10 | PSMB10 | 16q22.1 |
| MB1 | PSMB5 | 14q11.2 |
| delta | PSMB6 | 17p13 |
| Z | PSMB7 | 9q34.11-q34.12 |
| LMP7 | PSMB8 | 6p21.3 |
| LMP2 | PSMB9 | 6p21.3 |
| TAP1 | TAP1 | 6p21.3 |
| TAP2 | TAP2 | 6p21.3 |
| Tapasin | TAPBP | 6p21.3 |

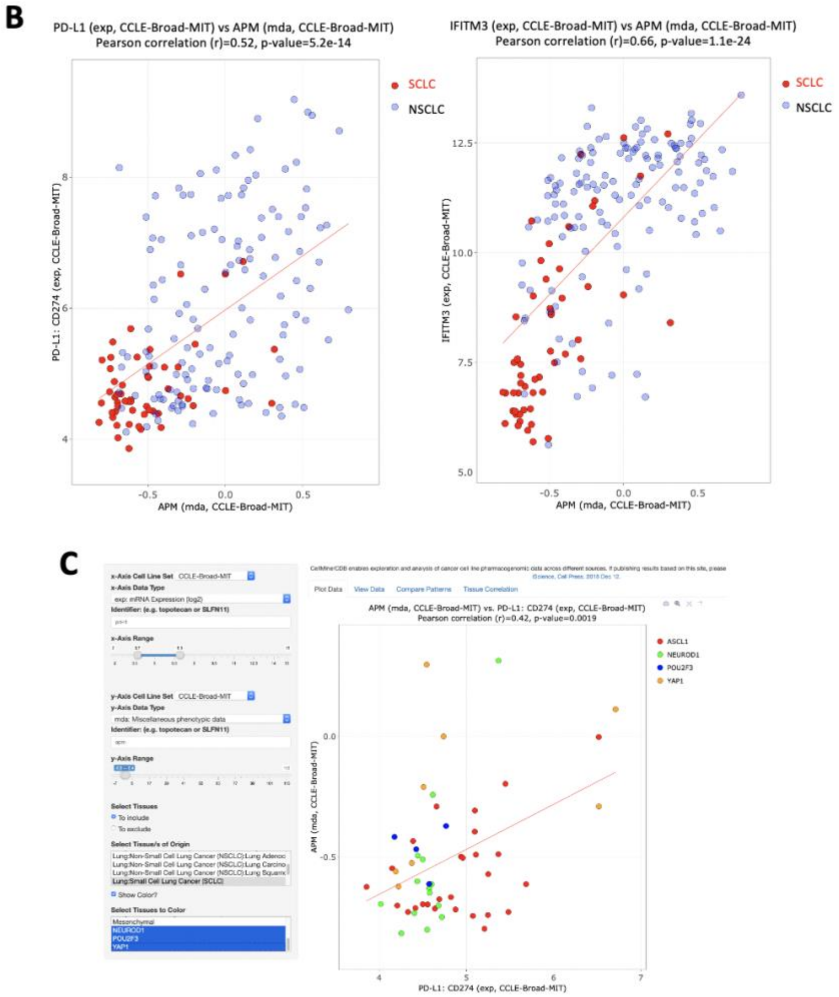


**Figure S13: Antigen presentation machinery (APM) signature in the SCLC cell lines**

(**A**) List of the 18 genes constituting the APM signature included as metadata in the CellMiner websites (https://discover.nci.nih.gov/CellMinerCDB and https://discover.nci.nih.gov/SclcCellMinerCDB). (**B**) Correlation between the APM score (x-axis) and the expression of *PDL1* (left) and *IFTIM3* (right). Each point represents a cell line (red: SCLC and blue: NSCLC cell lines). Note the lower AMP score in the SLCL compared to the NSCLC cell lines. (**C**) *PD-L1* expression (x-axis) vs the AMP score (y-axis) in the 116 cell lines of SCLC_CellMiner_Global. Note that only few cell lines have high AMP score. Among the cell lines with highest AMP score, most of them are SCLC-Y. Each point represents a cell line (red: SCLC-A, green: SCLC-N, blue: SCLC-P and orange: SCLC-Y cell lines).

#### SCLC_CellMiner: Integrated Genomics and Therapeutics Predictors of Small Cell Lung Cancer Cell Lines based on their genomic signatures

Camille Tlemsani, Lorinc Pongor, Luc Girard, Nitin Roper, Fathi Elloumi, Sudhir Varma, Augustin Luna, Vinodh N. Rajapakse, Sebastian Robin, Kurt W. Kohn, Julia Krushkal, Beverly A. Teicher, Paul S. Meltzer, William C. Reinhold, John D. Minna, Anish Thomas and Yves Pommier

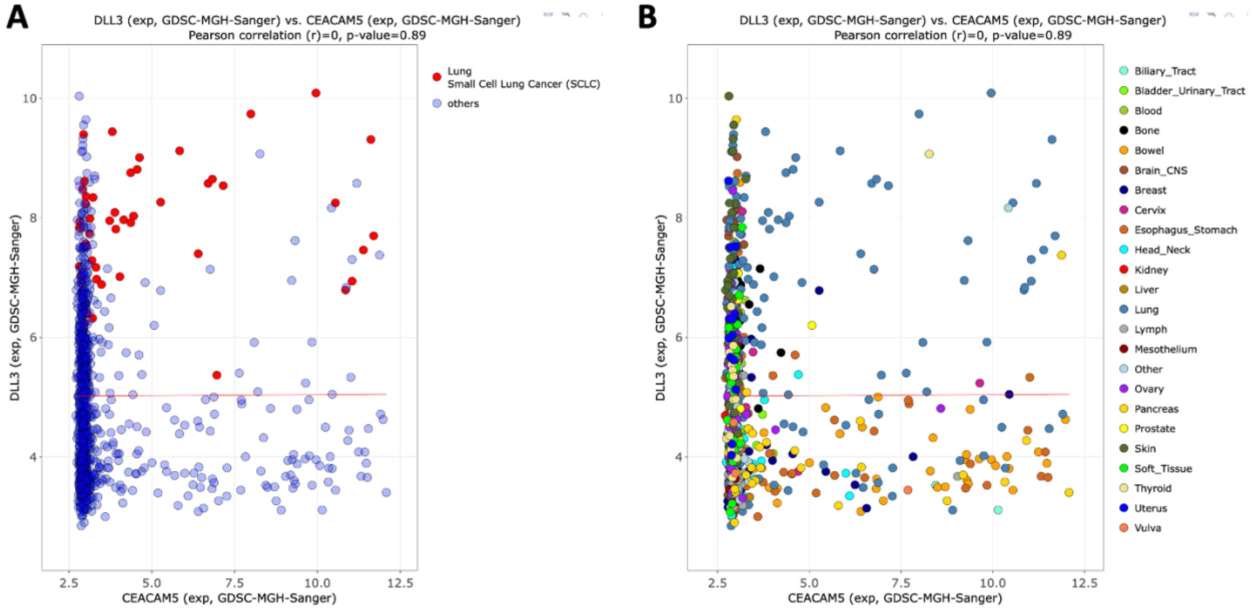


**Figure S14: *CEACAM5* is expressed in SCLC but not correlated with *DLL3* expression**

Snapshot of CellMinerCDB (https://discover.nci.nih.gov/cellminercdb) showing *CEACAM5* expression (x-axis) vs *DDL3* expression (y-axis) in the 986 cell lines of the GDSC dataset. Each point represents a cell line. (**A**) SCLC are in red and the other cell lines in blue. Most SCLC cell lines have a high expression of *DLL3* but only a subset has high expression of *CEACAM5*. (**B**) Cell lines are represented according to their histological subtype. SCLC cell lines (light blue) have both high *DLL3* and *CEACAM5* expression.

**SCLC_CellMiner: Integrated Genomics and Therapeutics Predictors of Small Cell Lung Cancer Cell Lines based on their genomic signatures**

Camille Tlemsani, Lorinc Pongor, Luc Girard, Nitin Roper, Fathi Elloumi, Sudhir Varma, Augustin Luna, Vinodh N. Rajapakse, Sebastian Robin, Kurt W. Kohn, Julia Krushkal, Beverly A. Teicher, Paul S. Meltzer, William C. Reinhold, John D. Minna, Anish Thomas and Yves Pommier

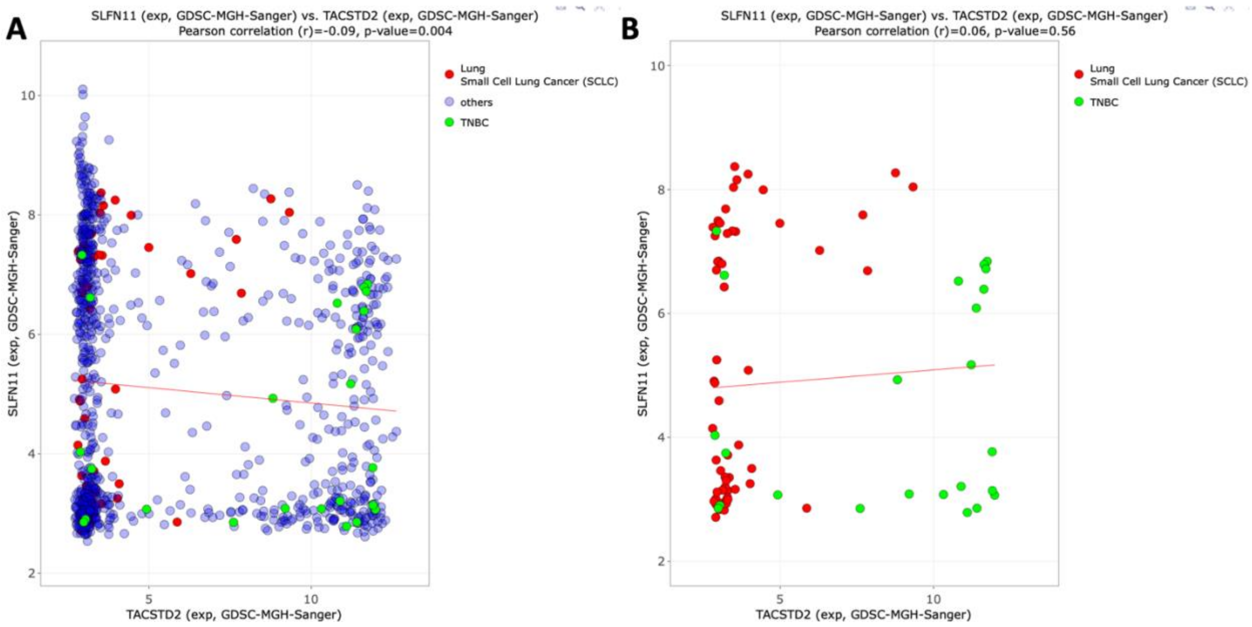


**Figure S15: Low expression of *TACSTD2* in the SCLC cell lines**

(**A**) Snapshot of CellMinerCDB (https://discover.nci.nih.gov/cellminercdb) showing the expression of *TACSTD2* (x-axis) vs *SLFN11* (y-axis) in the 986 cell lines of the GDSC dataset and in the subset of 59 SCLC and 24 triple negative breast cancer (TNBC) cell lines of the GDSC dataset (**B**). Each point represents a cell line (red: SCLC, green: TNBC and blue: other cell lines).

**SCLC_CellMiner: Integrated Genomics and Therapeutics Predictors of Small Cell Lung Cancer Cell Lines based on their genomic signatures**

Camille Tlemsani, Lorinc Pongor, Luc Girard, Nitin Roper, Fathi Elloumi, Sudhir Varma,
Augustin Luna, Vinodh N. Rajapakse, Sebastian Robin, Kurt W. Kohn, Julia Krushkal, Beverly A. Teicher,
Paul S. Meltzer, William C. Reinhold, John D. Minna, Anish Thomas and Yves Pommier

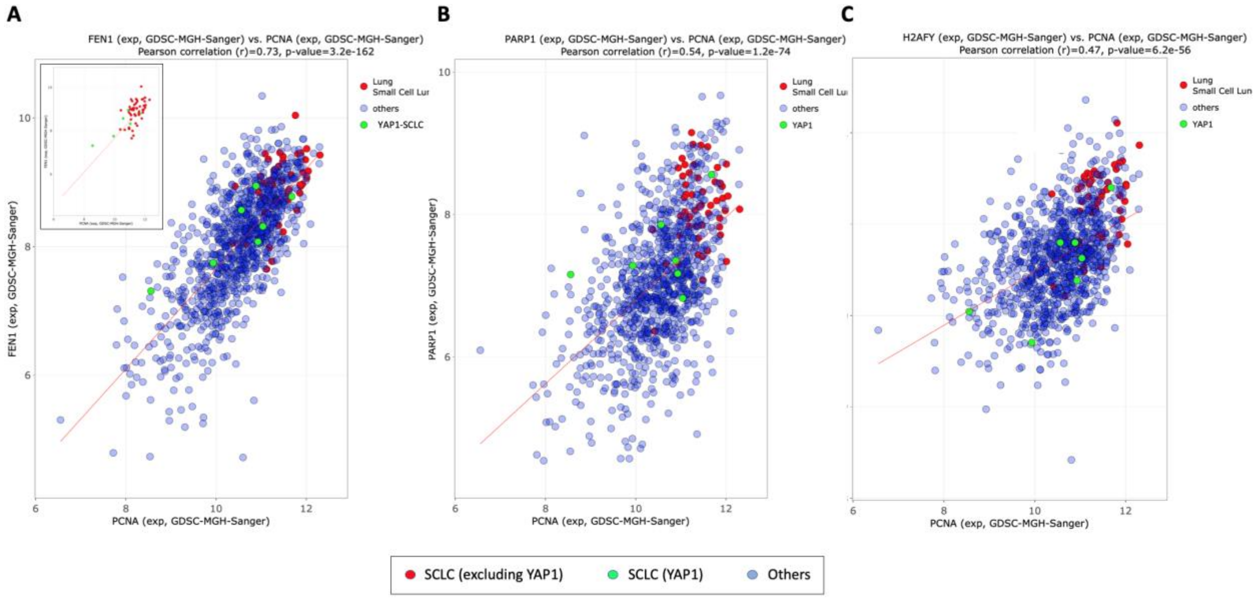


#### Figure S16: Overexpression of selected DNA replication genes in the SCLC cell lines

Snapshot of CellMinerCDB (https://discover.nci.nih.gov/cellminercdb) showing the expression of *PCNA* (x-axis) vs *FEN1* (**A**), *PARP1* (**B**) and *H2AFY* (**C**) (y-axis) in all histological subtypes of cell lines from GDSC dataset. Each point represents a cell line (red: SCLC, green: SCLC-Y cell lines and blue: other cell line subtypes). Note that DNA replication genes are highly expressed in SCLC cell lines except for the SCLC-Y cell lines.

#### SCLC_CellMiner: Integrated Genomics and Therapeutics Predictors of Small Cell Lung Cancer Cell Lines based on their genomic signatures

Camille Tlemsani, Lorinc Pongor, Luc Girard, Nitin Roper, Fathi Elloumi, Sudhir Varma,
Augustin Luna, Vinodh N. Rajapakse, Sebastian Robin, Kurt W. Kohn, Julia Krushkal, Beverly A. Teicher,
Paul S. Meltzer, William C. Reinhold, John D. Minna, Anish Thomas and Yves Pommier

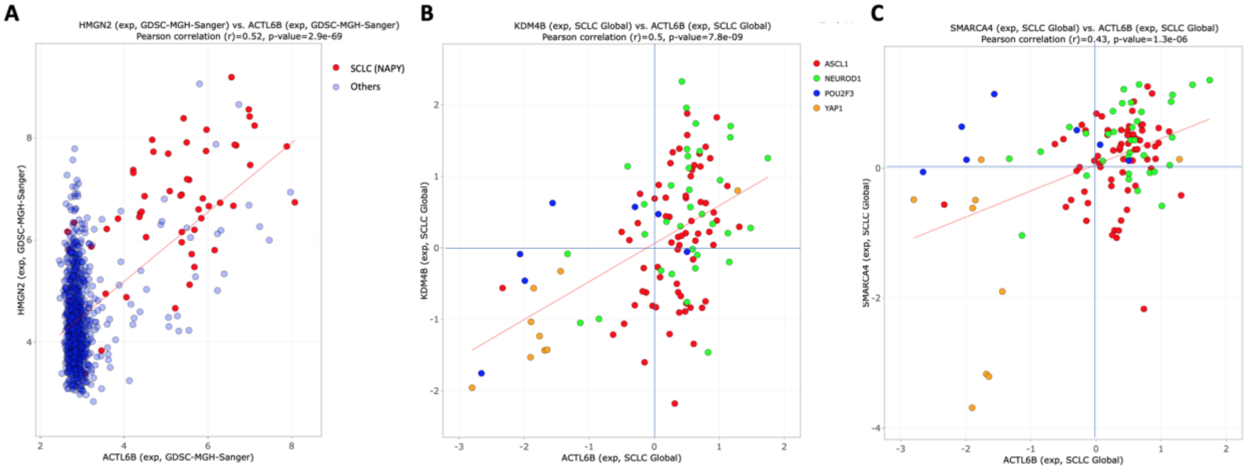


**Figure S17: High expression of *ACTL6B* is specific to the neuroendocrine SCLC cell lines**

(**A**) Snapshot of CellMinerCDB (https://discover.nci.nih.gov/cellminercdb) showing both high *ACTL6B* (x-axis) and *HMGN2* expression (y-axis) in the 59 SCLC cell lines (red) of the 986 GDSC cell lines. (**B-C**) Snapshot of SCLC_CellMiner (https://discover.nci.nih.gov/SclcCellMinerCDB) showing co-expression of *ACTL6B* (x-axis), *KDM4B* (**B**) and *SMARCA4* (**C**) (y-axis) among the 116 cell lines of SCLC_Global (green: SCLC-N, red: SCLC-A, blue: SCLC-P and orange: SCLC-Y cell lines). Note that among the SCLC cell lines, most of the cell lines expressing low *ACTL6B* are non-neuroendocrine (SCLC-P and SCLC-Y).
